# Successful intergeneric transfer of a major apple scab resistance gene (*Rvi6*) from apple to pear and precise comparison of the downstream molecular mechanisms of this resistance in both species

**DOI:** 10.1101/2021.05.31.446424

**Authors:** L. Perchepied, E. Chevreau, E. Ravon, S. Gaillard, S. Pelletier, M. Bahut, P. Berthelot, R. Cournol, H.J. Schouten, E. Vergne

**Author notes:** Corresponding author E. Vergne. L. Perchepied and E. Chevreau made equal contributions to this work.

## Abstract

**Background:** Scab is the most important fungal disease of apple and pear. Apple (*Malus x domestica* Borkh.) and European pear (*Pyrus communis* L.) are genetically related but they are hosts of two different fungal species: *Venturia inaequalis* for apple and *V. pyrina* for European pear. The apple/*V. inaequalis* pathosystem is quite well known, whereas knowledge about the pear/*V. pyrina* pathosystem is still limited. The aim of our study was to analyse the mode of action of a major resistance gene of apple (*Rvi6*) in transgenic apple and pear plants interacting with the two scab species (*V. inaequalis* and *V. pyrina*), in order to determine the degree of functional transferability between the two pathosystems.

**Results:** Transgenic pear clones constitutively expressing the *Rvi6* gene from apple were compared to a scab transgenic apple clone carrying the same construct. After inoculation in greenhouse with *V. pyrina*, strong defense reactions and very limited sporulation were observed on all transgenic pear clones tested. Microscopic observations revealed frequent aborted conidiophores in the *Rvi6* transgenic pear / *V. pyrina* interaction. The macro- and microscopic observations were very comparable to the *Rvi6* apple / *V. inaequalis* interaction. However, this resistance in pear proved variable according to the strain of *V. pyrina*, and one of the strains tested overcame the resistance of most of the transgenic pear clones. Comparative transcriptomic analyses of apple and pear resistant interactions with *V. inaequalis* and *V. pyrina*, respectively, revealed different cascades of molecular mechanisms downstream of the pathogen recognition by *Rvi6* in the two species. Signal transduction was triggered in both species with calcium (and G-proteins in pear) and interconnected hormonal signaling (jasmonic acid in pear, auxins in apple and brassinosteroids in both species), without involvement of salicylic acid. This led to the induction of defense responses such as a remodeling of primary and secondary cell wall, lipids biosynthesis (galactolipids in apple and cutin and cuticular waxes in pear), systemic acquired resistance signal generation (in apple) or perception in distal tissues (in pear), and the biosynthesis of phenylpropanoids (flavonoids in apple but also lignin in pear).

**Conclusion:** This study is the first example of a successful intergeneric transfer of a resistance gene among *Rosaceae*, with a resistance gene functioning towards another species of pathogen.

## Background

Apple (*Malus domestica* Borkh.) and European pear (*Pyrus communis* L.) are two closely related species of great economic importance for fruit production. A range of pests and diseases attacks both species and their production requires a high number of treatments. Scab, caused by *Venturia* species, is the most damaging fungal disease of both fruit species in all temperate countries. This disease causes necrotic lesions on leaves and fruits, which decrease the tree vigor and strongly reduce fruit quality, which make fruits unsuitable for fresh market sales. Chemical scab control under oceanic climates usually requires spraying up to 20 treatments per year and the development of alternative production systems (integrated protection, organic farming) reduce only partially the number of treatments [1]. A sustainable approach is the breeding of new varieties carrying durable resistance toward this disease. To achieve this goal, a better understanding of the function of major resistance genes and downstream defenses is needed.

Apple and European pears are hosts of two different fungal species: *Venturia inaequalis* for apple and *V. pyrina* (formerly named *V. pirina* [2]) for European pear. A long history of association between host and pathogen permitted their coevolution, which led to a narrow host spectrum for each *Venturia* species (i.e. genus specific) [3]. The level of genetic knowledge of the two pathosystems is very different. In the case of apple scab, numerous major resistance genes (R genes) and quantitative trait loci (QTLs) have been identified [4] and apple/*V. inaequalis* was one of the first plant pathosystem with good evidence for gene-for-gene interactions [5]. On the contrary, knowledge about the pear/*V. pyrina* pathosystem is still limited. *V. pyrina* presents at least five physiological races, which were found to have a very narrow range of pathogenicity [6]. So far, only one R gene and several QTLs have been identified [7].

*Rvi6* (formerly *HcrVf2*) is a major scab R gene which has been widely used in apple breeding programs. It is the first resistance gene of apple which has been isolated [8]. It is a receptor-like protein (RLP) gene containing an extracellular leucine-rich repeat and a putative transmembrane domain, resembling those of the *Cf9* tomato gene of *Cladosporium fulvum* resistance [9]. Transgenic apple lines expressing *Rvi6* under various promoter sequences present a strong resistance to several Rvi6-avirulent scab strains [10]. Genome-wide molecular analyses of the plant responses to scab have rarely been performed and only on apple host interactions. Subtractive hybridization [11, 12] and cDNA-AFLP [13] led to the identification of a limited set of differentially expressed genes in *Rvi6* natural resistant ‘Florina’ variety (scab inoculated ‘Florina’ versus mock, [12]), or in *Rvi6* resistant transgenic ‘Gala’ lines (*Rvi6* transgenic ‘Gala’ versus non-transformed ‘Gala’, after scab inoculation, [11]; *Rvi6* transgenic ‘Gala’ before versus post scab inoculation, [13]). RNA-seq identified five candidate genes putatively involved in the ontogenic scab resistance of apple [14]. In addition, nuclear proteome analysis identified 13 proteins with differential expression patterns among varying scab resistance ‘Antonovka’ accessions [15]. Therefore, in-depth knowledge of transcriptional patterns and gene functions involved in apple and pear scab resistance is still needed.

Plant immune receptors are the initial key step for recognition of invading pathogens and signalization of plant efficient defense mechanisms. Engineering plants via transfer of such R genes has the potential to increase disease resistance in many crops. Many R genes have now been shown to maintain their function after transfer to other plant species (reviewed in [16, 17]). In most cases, these transfers proved successful inside the *Solaneaceae* or the *Poaceae* families but efficient transfers have also been obtained interfamily or even across the monocot and dicot clades. Several classes of R genes have been successfully transferred between plant species: receptor kinase (RK), RLP, nucleotide binding leucine-rich repeat (NLR). Transfer of R genes acting in a gene-for-gene manner to another pathosystem implies two elements: 1) similarity of pathogen effectors recognized by the R gene and 2) sufficient conservation of downstream signaling pathway leading to efficient defense responses. To our knowledge, transfer of R genes between different pathosystems in the Rosaceae family has never been reported.

The objectives of our study were: 1) the functional transfer of the apple scab *Rvi6* gene to the pear/*V. pyrina* pathosystem, 2) the molecular dissection of *Rvi6*-mediated defense responses in the two pathosystems.

## Results and discussion

### Efficient production of pear transgenic lines

The transgenic clone P_MdRbc_HcrfVf2-11 (GalaRvi6) used in this study was already described by Joshi et al. [10] and get from Wageningen UR Plant Breeding.

The binary plasmid pMF1-pMdRbc1.6-Vf2-tMdRbc [10] containing the *Rvi6* coding sequence and the regulatory promoter and terminator sequences from the apple *Rubisco* gene was used to transform the ‘conference’ pear variety. In total, 78 kanamycin resistant lines were produced in a single transformation experiment on 650 ‘Conference’ leaf explants, reaching a rate of transformation of 12%. This rate is in the higher range of efficiency of most reports of pear transformation [7]. A sample of 30 lines was checked for ploidy level by flow cytometry. Three tetraploid lines and one chimeric (2n/4n) line were discarded. Polymerase chain reaction (PCR) analysis confirmed the presence of the *Rvi6* transgene and the absence of *Agrobacterium* contamination in the remaining 26 lines. A sample of eleven transgenic lines were rooted and acclimatized for scab inoculation in greenhouse.

Quantitative polymerase chain reaction (Q-PCR) analyses were performed on leaf samples of these lines at two separate times (spring and autumn) to evaluate the level of expression of the transgene *Rvi6*. Results in Fig. 1 indicate a large variability of relative expression of the transgene among the transgenic lines, in the range of 50 to 500 compared to a background level in ‘Conference’, which does not possess *Rvi6*. Similarly, Joshi et al. [10] observed a wide variation of expression among apple transgenic lines expressing *Rvi6* controlled by the same apple small subunit rubisco gene promoter (*MdRbc*) (57-163 compared to the natural expression level of this resistance gene in the apple cultivar ‘Santana’ that obtained the *Rvi6* gene by means of conventional breeding), that was not correlated with the copy number of the transgene. In our results on pear, the expression levels measured in spring were generally lower than those measured in autumn, but a consistent ranking of the lines was obtained in the two assays. The clone 60AU appeared as the highest expressor of *Rvi6* in this sample. Therefore, this line was chosen for the subsequent transcriptomic analyses.

**Fig. 1:**
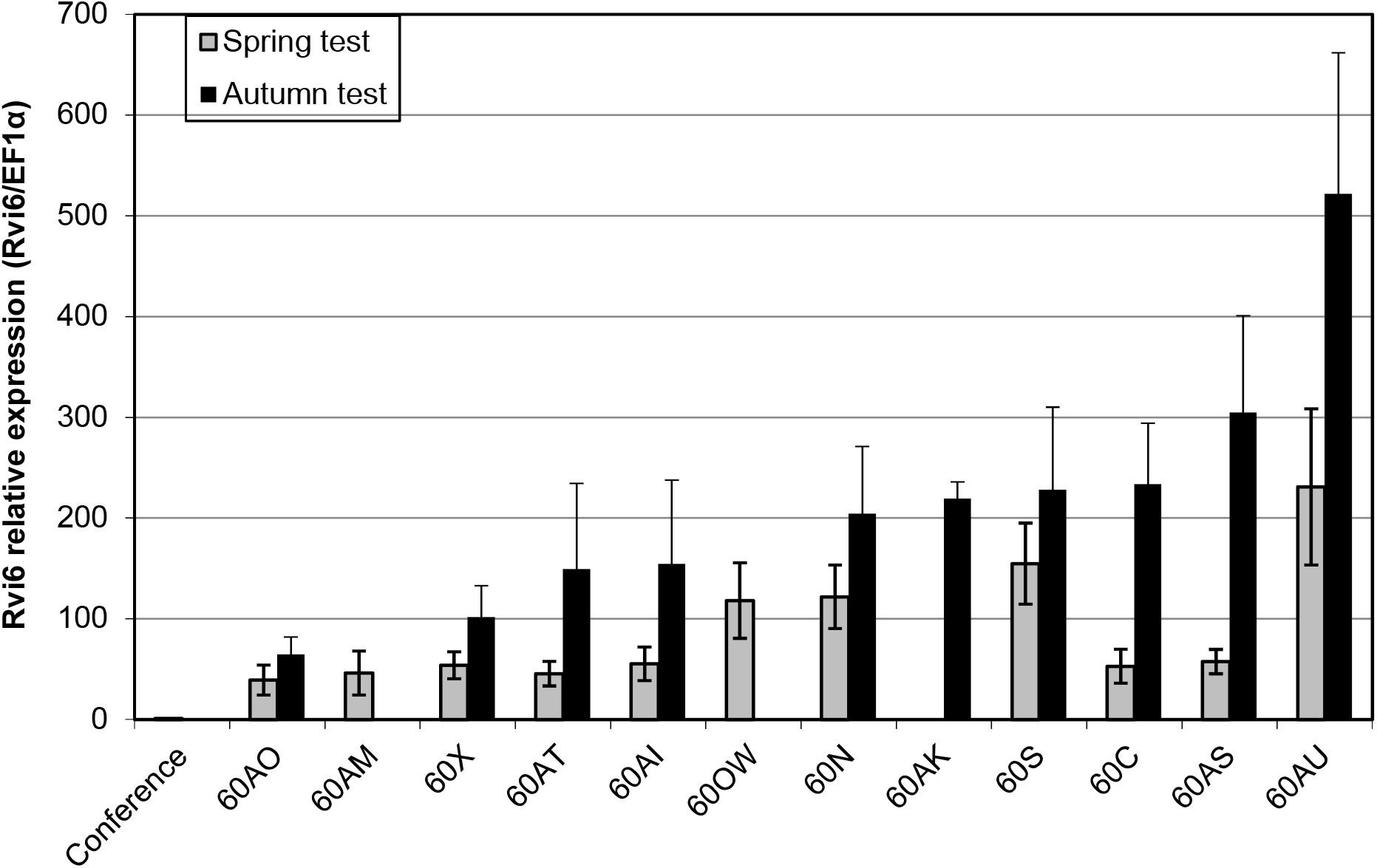
Level of expression of the transgene *Rvi6* in transgenic pear lines. Bars are the mean of three independent replicates, error bars indicate confidence intervals at α=0.05. Normalization was done with the reference gene *EF1α* and the non-transgenic genotype ‘Conference’ was used as a calibrator.

### High level of *V. pyrina* resistance in pear transgenic lines but possible breakdown with some *V. pyrina strains*

A scab inoculation test was performed on nine independent pear transgenic lines carrying the *Rvi6* transgene. The area under the disease progress curve (AUDPC) based on sporulation scores at 14, 21, 28, 35 and 42 days after inoculation summarizes the results (Fig. 2). The three strains of *V. pyrina* caused typical severe scab (100% of class 4 symptoms, Table S1) on the non-transgenic ‘Conference’ (susceptible control). Very strong resistance was observed in the nine transgenic lines challenged with strain VP 137, with only 2% of the plants in susceptible class 3b. All the tested transgenic lines were also clearly resistant to strain VP 102, with only 6% of the plants in susceptible class 3b. However, various levels of susceptibility were observed among transgenic pear lines inoculated with strain VP 98. In total, 57 % of the plants from all transgenic lines produced susceptible symptoms of classes 3b or 4. Even though the AUDPC of all transgenic lines was significantly lower than ‘Conference’, this indicates a partial breakdown of *Rvi6* resistance in several transgenic pear lines. These results are very similar to the results of Joshi et al. [10] who tested the scab resistance of apple transgenic lines expressing the same construct. They observed total resistance of the transgenic apple lines towards four isolates of *V. inaequalis* avirulent on *Rvi6*-based resistant cultivars, but this resistance was overcome by the isolate EU-D42, virulent on *Rvi6*-based resistant cultivars. Little is known so far about the effectors of *Venturia* species and the basis of their host specificity. Whole genome sequencing allowed a comparative analysis of the predicted secretomes of *V. inaequalis* and *V. pyrina* [18]. This led to the identification of many candidate effector genes or gene families, some of them being unique to *V. inaequalis* or to *V. pyrina* isolates. Recently, the *AvrRvi6* from *V. inaequalis* has been identified as a 93 amino acid protein containing 6 cysteines [19], but no precise homologous has yet been identified in *V. pyrina*. Our findings indicate that the apple transgene *Rvi6* expressed in pear probably recognizes avirulence effectors similar to *AvrRvi6*, secreted by *V. pyrina*, and that some *V. pyrina* strains possess virulence factors leading to *Rvi6* resistance breakdown. No significant correlation could be found among the transgenic pear lines between the expression level of *Rvi6* and the degree of resistance or the class of symptoms. The *MdRbc* promoter of apple that was used in this study proved to give high levels of expression in apple [20]. Therefore, we can assume that the levels of expression of *Rvi6* in the transgenic pear lines were all above the necessary threshold to induce a strong resistance reaction. Similarly, Joshi et al. [10] observed that expression of *Rvi6* beyond a certain level did not result in increased resistance of the transgenic *Rvi6* apples lines nor in an extension of the spectrum of resistance towards virulent strains.

**Fig. 2:**
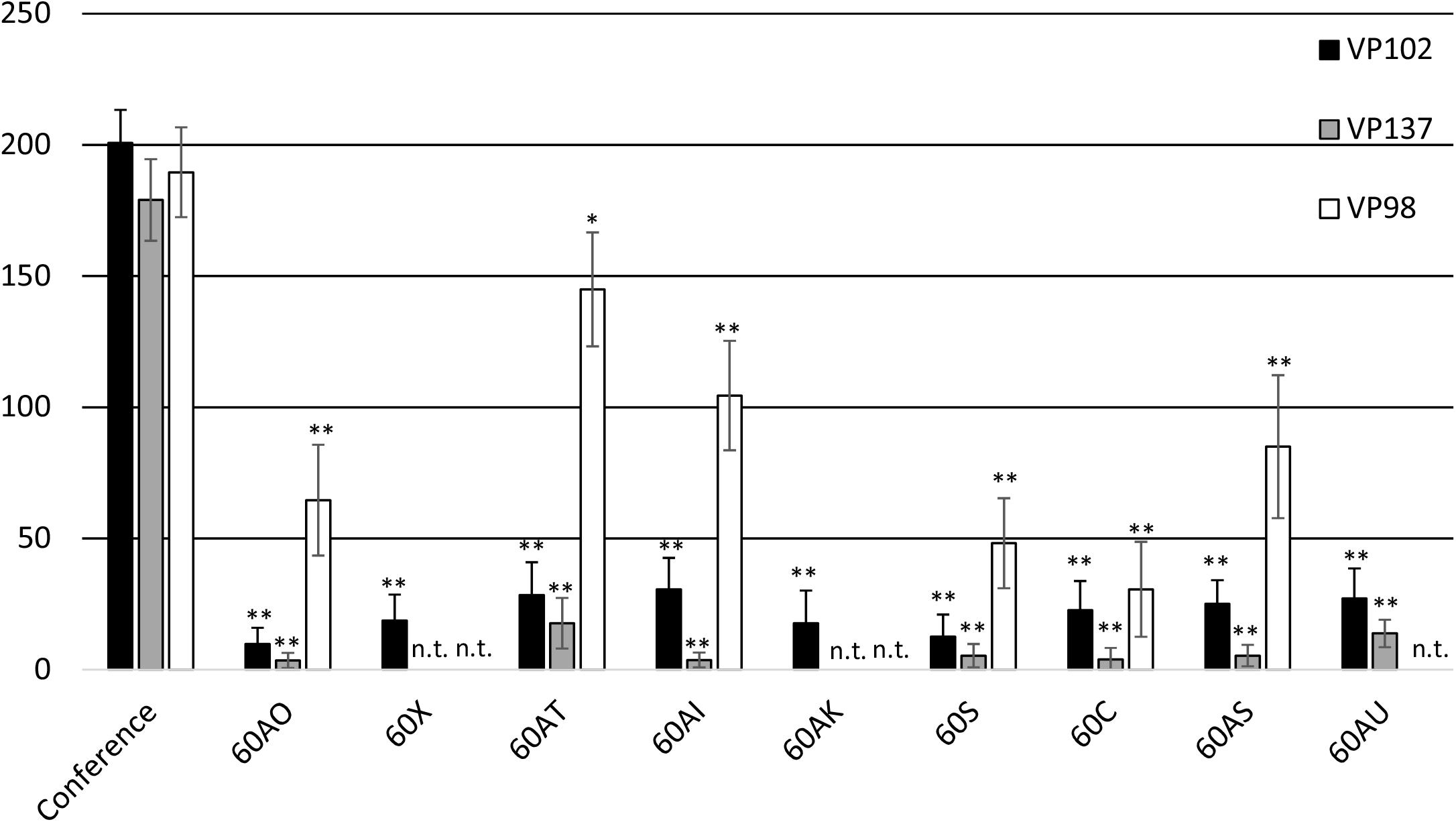
Scab susceptibility of transgenic pear lines inoculated with three different *V. pyrina* strains. AUDPC based on sporulation scores at 14, 21, 28, 35 and 42 days after inoculation with three different *V. pyrina* strains (VP98, VP102, VP137) on a series of transgenic Conference genotypes that received the *Rvi6* gene from apple by means of stable transformation. Bars are the mean of 13 to 25 shoots, error bars indicate confidence intervals at α=0.05.

### Variable expression of resistance symptoms

At the macroscopic level (Fig. 3A), susceptible interactions (apple: Gala/*V. inaequalis* and pear: Conference/*V. pyrina*) led to a strong sporulation appearing on the upper side of the leaves, but also in the case of pear on the lower side of the leaves as well as on the shoots, as observed previously on ‘Conference’ [21]. At the microscopic level, the kinetics of fungal development was very similar in apple and pear susceptible interactions. Conidia germination and formation of appressoria were achieved three days after inoculation (Fig. 3B). Seven days after inoculation, numerous conidiophores were formed and released conidia (Fig. 3C, 3D), the number of scars indicating the intensity of sporulation (Fig. 3E).

**Fig. 3:**
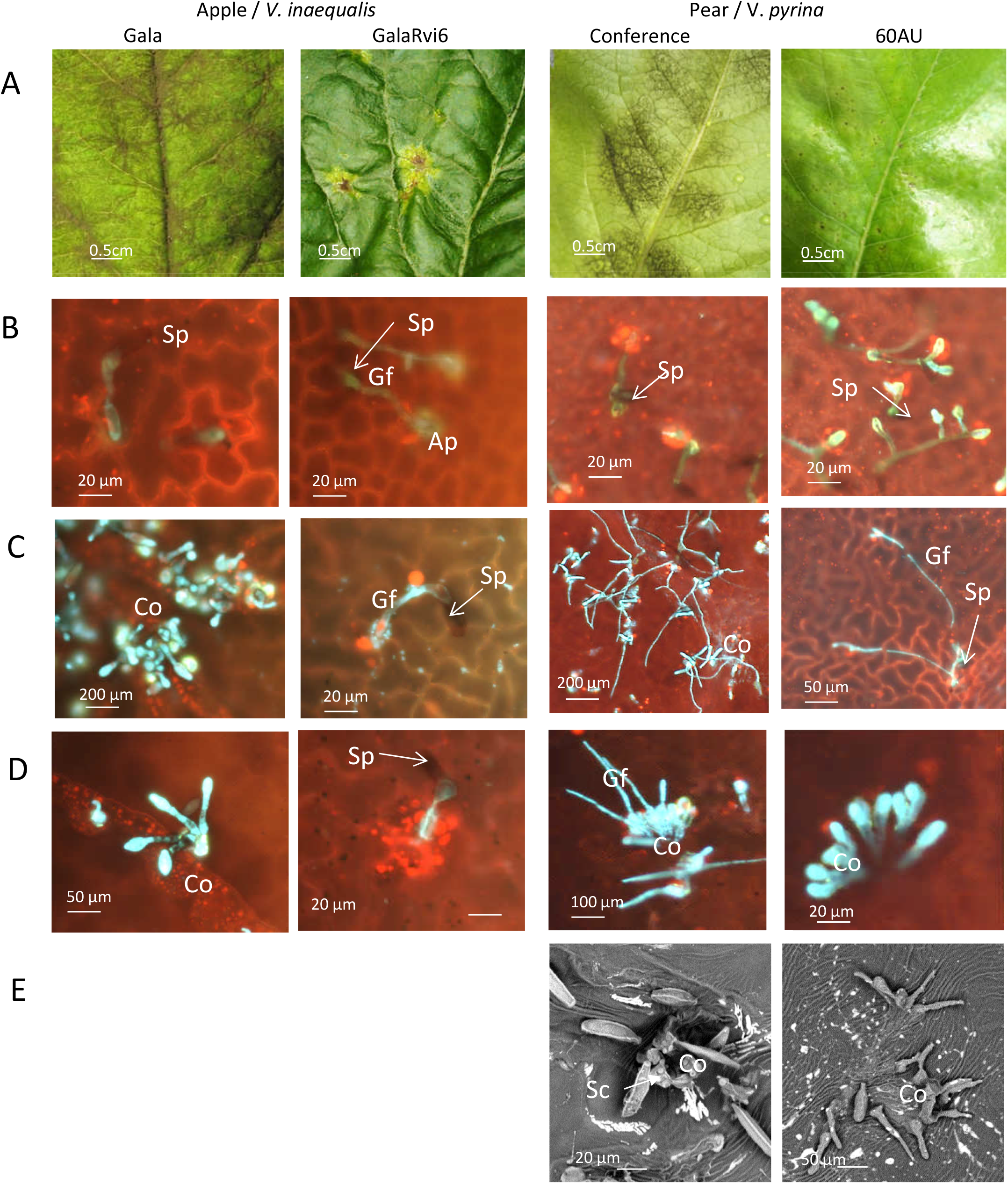
Macro- and microscopic observations of apple and pear susceptible and resistant interactions. (A) binocular observations 42 days after inoculation; (B) wide field fluorescence observations 3 days after inoculation; (C) wide field fluorescence observations 7 days after inoculation (D) wide field fluorescence observations 14 days after inoculation; (E) SEM observation 19 days after inoculation. Ap: appressorium, C: conidia, Co: conidiophore, Gf: germination filament, Sc: scar, Sp: spore.

At the macroscopic level (Fig. 3A), resistant interactions (GalaRvi6/*V. inaequalis* and 60AU/*V. pyrina*) produced very variable symptoms. In addition to pinpoints, chlorotic and necrotic lesions, with or without leaf crinkling, were frequently observed in apple as well as in pear. At the microscopic level, apple resistance was characterized by short germination filaments. In addition, infected sites surrounded by a ring of red autofluorescent cells (probably due to accumulation of phenolic compounds) around the appressoria were frequently observed. Subcuticular stroma was visible, but no conidiophores were observed (Fig. 3B, C and D). In pear, long branched germination filaments were frequent (Fig. 3C) and many aborted conidiophores without conidia emission were observed (Fig. 3D and E).

The kinetics of establishment of the susceptible interactions is in agreement with previous reports on *V. inaequalis* [22, 23] and *V. pyrina* [21]. Similarly, the large range of resistance symptoms, from pin-points typical of hypersensitive reaction (HR) to chlorotic lesions with occasional very slight sporulation, has been frequently observed in apple genotypes carrying the *Rvi6* gene, get by conventional breeding [24] as well as on pear cultivars carrying partial to strong resistance [25]. The microscopic observations fit with the histological description of resistance symptoms of class 1, 2 or 3a in *Rvi6* apple genotypes [22].

Based on these findings, we decided to sample for the transcriptomic study at 8, 24 and 72 hours post inoculation (hpi), in order to cover the period of establishment of the first intimate contact between fungal and plant cells.

### Common and specific patterns of gene expression modulation during the first steps of *Rvi6*-induced resistance in apple versus pear

Understanding the intricate mechanisms of *Rvi6*-induced resistance in apple and pear required the genome-wide analysis of differentially expressed genes between susceptible non-transgenic and resistant *Rvi6* expressing lines, and their comparison between the two species. Several studies reported differential apple transcriptomic responses to different biotic stresses, but most of the studied pathogens were necrotrophs, with the exception of Tao et al. [26] who studied the compatible apple interaction with *Gymnosporangium yamadae*. A recent meta-analysis of these studies showed that cell wall genes of apple were downregulated in all three datasets: fungal pathogens (*Valsa mali, Alternaria alternata, Marssonina coronaria, Pythium ultimum, Venturia inaequalis*), virus (Apple Stem Grooving Virus) and bacteria (*Erwinia amylovora*) [27]. Concerning the fungal pathogens, several genes involved in flavonoids (both chalcones and dihydroflavonols) and isoprenoids mevalonate pathways were upregulated. PR proteins and other stress related proteins were also upregulated. Fungal pathogens predominantly upregulated the genes involved in ethylene, brassinosteroids (BRs) and salicylic acid while jasmonic acid (JA) responses were mostly repressed. Ubiquitin-mediated degradation was mainly upregulated. Fungal pathogens also induced UDP glycosyltransferases, phosphatases, nitrilase and glutathione-s-transferase. More precisely, two studies concerned *Rvi6*-mediated resistance and identified a limited number of differentially expressed genes (DEGs). For example, Paris et al. [11] showed that protein degradation genes accounted for 13% of expressed sequence tags in *HcrVf2*-transformed apple plants in response to *V. inaequalis*. Cova et al. [12] showed that cinnamate 4-Hydroxylase involved in phenypropanoid pathway was activated in a *Rvi6* variety in response to *V. inaequalis* and after salicylic acid (SA) treatment. Scab ontogenic resistance of ‘Golden delicious’ was studied by Gusberti et al. [14] by RNA sequencing (RNA-Seq) analysis. They showed that EDS1, PAL and CAD genes were repressed while LOX was up-regulated during *V. inaequalis* infection. Therefore, the understanding of the molecular mechanisms underlying apple scab resistance are still very limited and, to our knowledge, no studies had been published on pear scab.

In our study, DEGs were analysed by comparing transcript abundance in leaves between susceptible non-transgenic and resistant *Rvi6* expressing lines, in apple and in pear, at each of the three time-points of the interaction with *V. inaequalis* and *V. pyrina* respectively. In total, 2977 DEGs in apple and 4170 DEGs in pear were identified, which amounts to 9.5 % of all apple genes on the apple “Aryane V2” microarray, and 9.5 % of all pear genes on the “pear V1” microarray. (Table 1).

**Table 1:** Number of differentially expressed genes (DEGs) identified at each of the three time points.

|  | GalaRvi6 versus Gala |  |  |  | 60AU versus Conference |  |  |  |
| --- | --- | --- | --- | --- | --- | --- | --- | --- |
|  | 0 hpi | 8 hpi | 24 hpi | 72 hpi | 0 hpi | 8 hpi | 24 hpi | 72 hpi |
| Total number of DEGs* | 173 | 1799 | 823 | 539 | 1115 | 2415 | 922 | 273 |
| DEGs in % of all genes on the microarray** | 0.55 | 5.75 | 2.63 | 1.72 | 2.54 | 5.50 | 2.10 | 0.62 |
| % of upregulated DEGs | 19.1 | 56.8 | 60.9 | 30.4 | 46.2 | 76.3 | 81.5 | 49.1 |
| % of downregulated DEGs | 80.9 | 43.2 | 39.1 | 69.6 | 53.8 | 23.7 | 18.5 | 50.9 |
| % of DEGs without TAIR name | 2.89 | 4.84 | 6.80 | 5.19 | 13.9 | 14.0 | 10.1 | 14.3 |

\*: DEGs numbers were calculated using the p-adj values $\leq 0.01$ as selection threshold
\*\* : 31311 genes on the apple Ariane V2 microarray, 43906 genes on the pear V1 microarray

In apple GalaRvi6 as in pear 60AU transgenic lines, *Rvi6* is under the control of the strong constitute Rubisco gene promoter. However, the reaction to these constitutive *Rvi6* expression is quite different between apple and pear in terms of DEG quantity. Indeed at T0, before scab inoculation, only 173 DEGs were detected between ‘Gala’ and GalaRvi6, among which 81 % were downregulated. On the contrary, in pear, 1115 DEGs between ‘Conference’ and 60AU were detected at T0, among which 74 % were specific of this constitutive state. 46 % of these DEGs were up-regulated. Using MapMan to map the DEGs TAIR names, we observed that beside protein and RNA metabolisms, the main functional categories represented in this set of DEGs were signaling, cell cycle, transport, stress and development (Fig. S1).

In both species, the greatest transcriptomic divergence between the susceptible and the resistant transgenic lines occurred at 8 hpi (with 1799 DEGs in apple and 2415 DEGs in pear), with 85 and 83 % of these DEGs specifically detected at 8 hpi and not later during the interaction, respectively. Among all the DEGs observed at 8, 24 and 72 hpi, the proportion of up-regulated DEGs was higher in pear (76 %) than in apple (53 %). The main functional categories represented in this set of DEGs were similar for both species: protein metabolism, RNA metabolism, signaling, transport and cell cycle (Fig. 4).

**Fig. 4:**
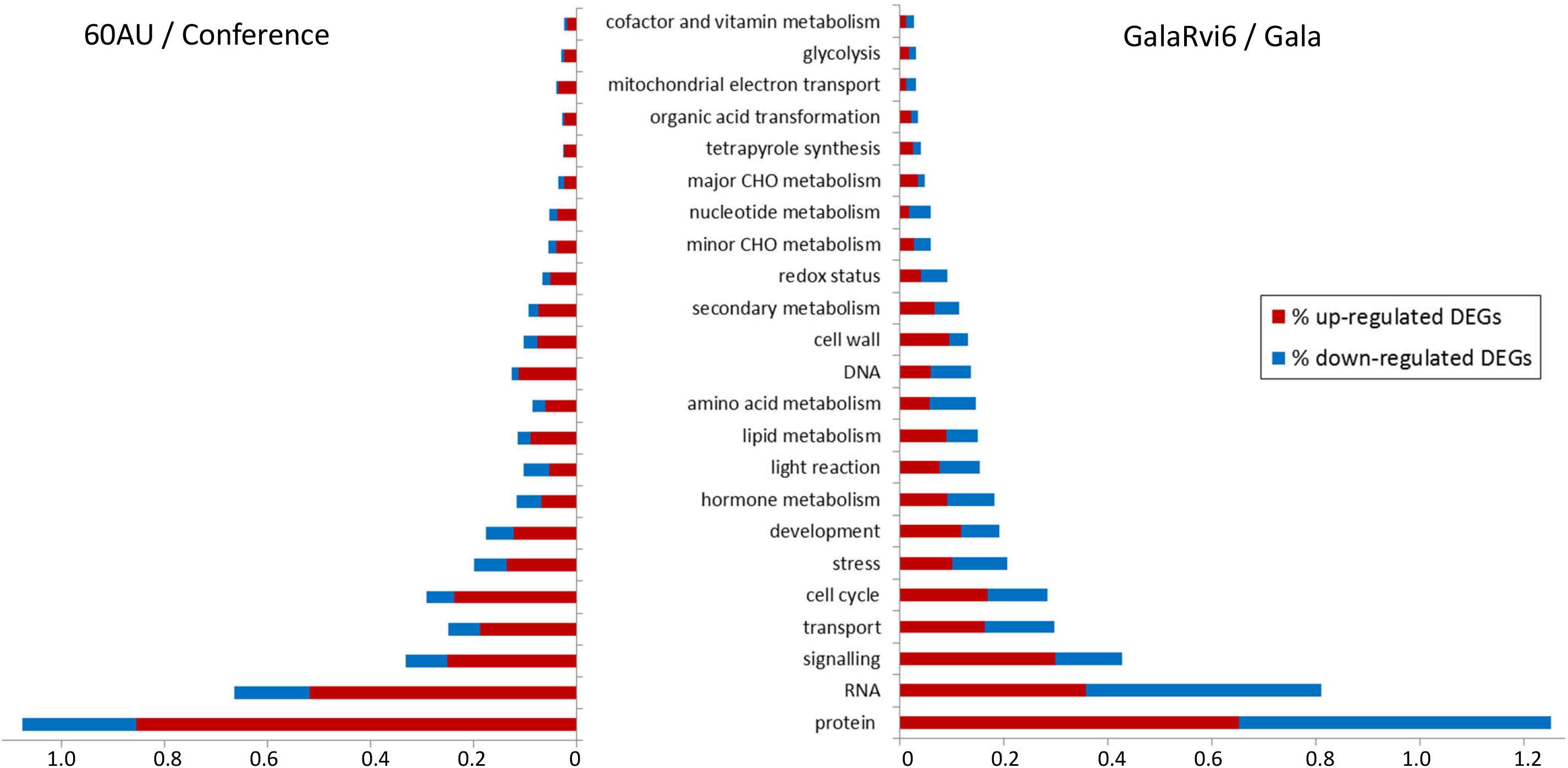
Functional categories of DEGs at 8, 24 or 72 hours post-inoculation. Pear (60AU / Conference, on the left) and apple (GalaRvi6 / Gala, on the right) responses to *V. pirina* and *V. inaequalis*. The number of up- or down-regulated DEGs is expressed as a percentage of the total number of genes present in the pear V1 (49306 genes) and Ariane V2 (31311 genes) microarrays, respectively. DEGs are classified in functional categories according to MapMan 3.5.1R2 bins. Only bins with ≥ 10 DEGs are presented.

To basically validate the transcriptomic data, 14 DEGS with varied ratios (between −2.01 and 3.57) have been tested in Q-PCR (Table S2), on the two biological repeats used for transcriptomic analyses. In this study 72 DEGs were discussed for apple, among them about 69% were at 8hpi, QPCR was thus focused on DEGs at that time. Concerning pear, 71 DEGs were discussed, among them about 89% were at 8 or 24hpi, and induced for a majority (66%). QPCR was then made essentially on DEGs with positive ratios at one of these two times (Table S2). The QPCR results confirmed the induced or repressed statute of tested DEGs.

Several functional categories of DEGs detected at 8, 24 or 72 hpi were further investigated.

#### Specific signaling receptors and pathways between apple and pear

In the signaling functional category, receptor like kinase (RLK), G-proteins and calcium related DEGs were predominant in apple and pear. Most of these genes were up-regulated as early as 8 hours post inoculation. Among the 33 and 30 receptor kinases up-regulated, 15 and 20 were RLK with a leucine rich repeat (LRR) domain in apple and pear respectively. Several of these RLKs could putatively be involved in effector-triggered immunity (ETI) or pattern-triggered immunity (PTI) (Fig. 5). For example, *CERK1*, which was up-regulated in apple, has a crucial role in glycan-based microbe-associated molecular pattern (MAMP) perception. CERK1 is implicated in chitin recognition and binding, although at low affinity. CERK1 has recently been shown to be necessary for 1,3-β-D-glucan-triggered immune responses. The 1,3-β-D-glucans are an important component of fungal and oomycete cell walls but are also present in plant cells, although in minute amounts, thus suggesting that CERK1 might also be a damage-associated molecular pattern (DAMP) receptor [28]. The central role of CERK1 in Arabidopsis immunity is supported by its role in resistance against fungi such as *Alternaria brassicicola, Golovinomyces cichoracearum* and *Plectosphaerella cucumerina*, the bacterium *Pseudomonas syringae* and the oomycete *Hyaloperonospora arabidopsidis* [29]. Among the RLKs, ATPERK1 (a proline extensin-like receptor kinase 1 gene) was up-regulated in apple. PERK1 shares sequence similarity to extensins and contains a transmembrane domain and a kinase domain. Silva and Goring [30] showed that PERK1 may be involved early on in the general perception and response to a wound and/or pathogen stimulus. Many pattern recognition receptors (PRR) form recognition complexes involving the multitask co-receptors BAK1 and SERK1 [31]. Both co-receptors were also up-regulated in apple. In contrast, a different array of receptors and co-receptors was found up-regulated in pear. Two negative regulators of BAK1 receptors complex formation (*BIR3* [32] and *ANX2* [33]) were up-regulated in pear. In contrast to apple, *CERK1* was down regulated in pear. Several DAMPs such as *RLK7* [34] were also up-regulated in pear. Thus *Rvi6*-mediated scab resistance seems to involve a different array of receptors and co-receptors in apple and pear.

**Fig. 5:**
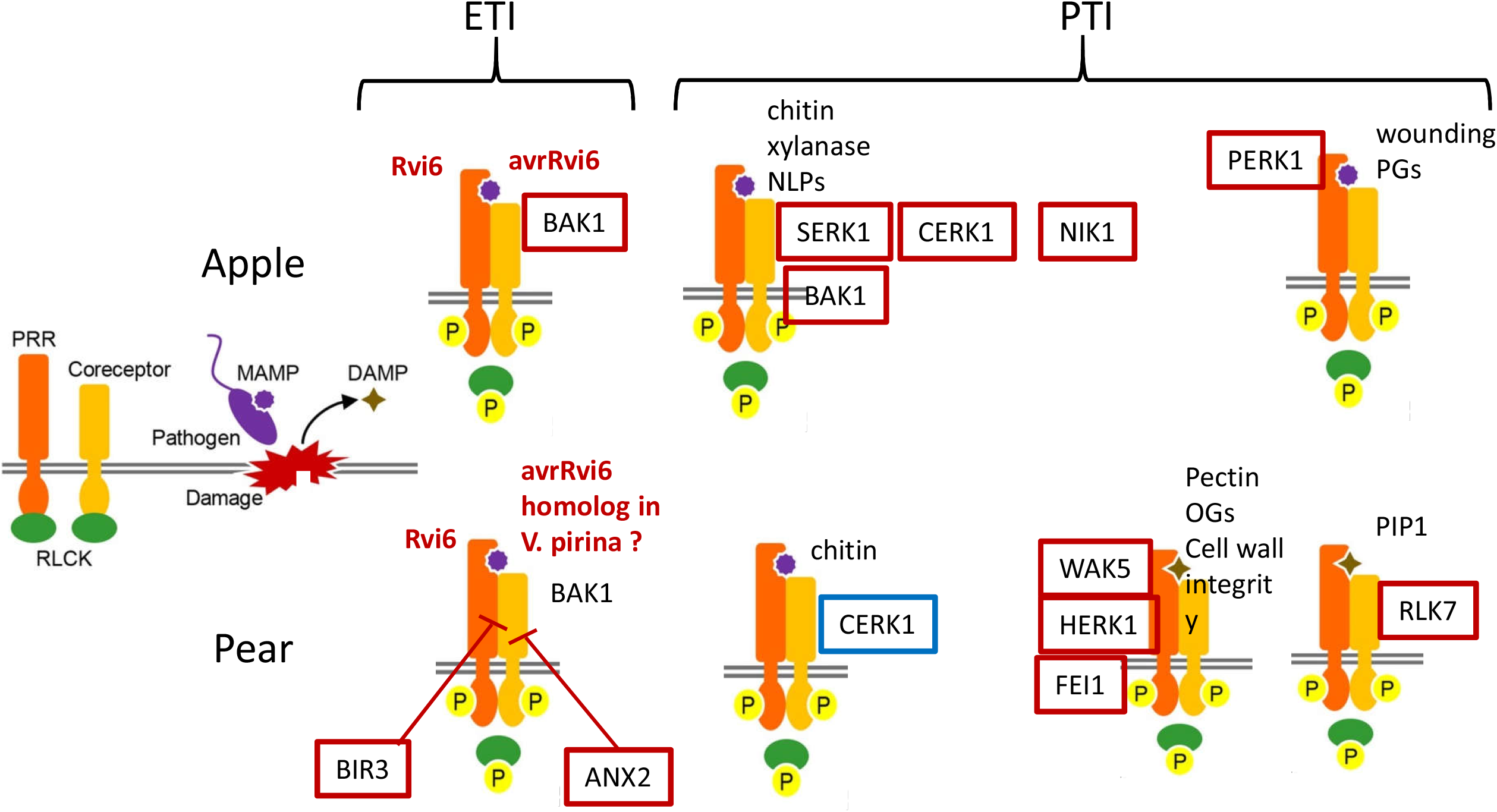
Main complexes of receptors and co-receptors putatively involved in ETI or PTI. Main related DEGs activated (in red) or repressed (in blue). Abbreviations: ANX2: ANXUR2; BAK1: BRI1-associated receptor kinase1; BIR3: BAK1-interacting LRR-RK3; CERK: LysM-RLK chitin receptor kinase; DAMP: damage-associated molecular pattern; HERK1 : HERCULES1; MAMP: microbe-associated-molecular pattern; NIK1: NSP-interacting kinase1; NLP: necrosis and ethylene-inducing peptide 1-like protein; OG : oligogalacturonides; PERK1: Proline Extensin-like Receptor kinase1; PG: polygalacturonase; PRR: pattern recognition receptor; RLCK: receptor-like cytoplasmic kinase; SERK1: somatic embryogenesis receptor kinase1; RLK7: Receptor-like kinase7; WAK5 : wall-associated kinase 5. (adapted from [117]).

Upon detection of specific pathogen effectors, NLR proteins mount a quick and robust reaction, usually culminating in a hypersensitive response to defend against and restrict further spread of the pathogen [35]. Because of the detrimental effects of R protein activation on plant cell growth and development, normally R protein-mediated immunity has to be under multiple levels of tight negative control [36]. The selective degradation of proteins largely occurs via the ubiquitin/26S proteasome pathway. In our results, genes involved in proteasome functions were mainly up-regulated in apple and pear, 62 out of 91 and 71 out of 98 DEGs respectively. Many DEGs up-regulated were ubiquitin-ligase E3s, 46 out of 56 in apple and 42 out of 61 in pear. Four main types of ubiquitin-ligase E3s are known in plants, and are classified by their mechanisms of action and subunit composition: HECT, RING, U-box and cullin–RING ligases (CRLs), with the CRLs further divided into four subtypes [37], including the S phase kinase-associated protein 1 (SKP1)–cullin 1 (CUL1)–F-box (SCF) E3s complex. In this last subtype, 13 out of 19 DEGs were up-regulated in pear and 9 out of 15 in apple. Members of the SCF (SKP1-CUL1-F-box protein) E3 ubiquitin ligase complex targets specific substrate proteins for ubiquitination and most often subsequent protein degradation. In our results, SKP1, CUL1 and CPR1 were up regulated and RPM1 was down regulated in pear. Overexpression of CPR1 abolishes immunity mediated by RPM1 and slightly compromises resistance specified by several other R proteins, such as RPP2 and RPP4, suggesting that SCF^CPR1^ may also target these R proteins for degradation [36]. These results are in favor of the hypothesis that NLR receptors do not take part in the HR development (pin-points) observed in the pear/*V. pyrina* interaction. SRFR1, up-regulated in apple, is another protein involved in the regulation of NLR protein-mediated immunity. SRFR1 functions in the negative regulation of NLR R protein accumulation [38]. Regulation by the SCF E3 ubiquitin ligase complex seems to be of great importance in apple and pear.

The main difference between apple and pear signaling DEGs concerned the G-protein signaling. In apple, only 14 G-protein DEGs were up-regulated (including 2 and 3 GTP-binding proteins of the RAB and ARF families respectively) and 14 DEGs were down-regulated (10 GTP-binding proteins of the RAB family out of 14). Most G-protein DEGs were up-regulated in pear (32 out of 36), including *NOG1* belonging to the family of TRAFAC (translation factors), 13 GTP-binding proteins of the RAB family and 2 GTP-binding proteins of the ARF family. Lee et al. [39] identified the small GTP-binding protein NOG1 as a novel player in guard cell signaling and early plant defense in response to bacterial pathogens. Heterotrimeric G-proteins are members of the superfamily of G-proteins. The G-protein hetero-trimer is composed of α, β, and γ subunits (G_α_, G_β_, and G_γ_), organized in a highly conserved structure, which is in complex with transmembrane G-protein-coupled receptors. Delgado-Cerezo et al. [40] suggested a canonical functionality of the G_β_ and G_γ1_/_γ2_ subunits in the control of Arabidopsis immune responses and the regulation of cell wall composition. Indeed the alteration in the content of xylose, such as in *det3* and *irx6* mutants, might explain the enhanced susceptibility to *Plectosphaerella cucumerina* observed in the G_β_ / G_γ_ mutants. *DET3* was up-regulated in apple and *IRX6* was down-regulated in pear (detailed later). Plant U-box type E3 ubiquitin ligases (PUBs) are implicated in the regulation of the immune response and cell death. PUB13 was also up-regulated in pear. Interestingly the fact that PUBs interact with two types of key signaling components, kinases and G-proteins, has emerged recently [41]. G-proteins act as molecular switches that are involved in transmitting signals from a variety of stimuli, maybe like presence of *V. inaequalis*.

Among the calcium related DEGs, calmodulin binding proteins (*IQD13* and *IQD31* in apple; *IQD6* in pear), calmodulin dependent protein kinase (CPK) (*CPK8* and *CPK28* in apple; *CPK1*, *CPK6*, *CPK13* and *CPK21* in pear) and calcium ion binding (*ATCP1, ATCBL3* in apple; *CAM3, CAM7, CRT3* in pear) were up-regulated. Thus calcium signaling appeared important in apple as well as in pear during the establishment of *Rvi6*-mediated scab resistance.

Among the hormonal pathways, most DEGs belonged to the auxin pathway in apple. Most of the genes involved in the auxin pathway were up-regulated. These DEGs were involved in biosynthesis (*JAR1* in apple), transport (*PIN1, ARG1, MDR1, AUX1* in apple; *PIN1* in pear), signaling (*ILR1* in apple) and regulation by auxin (*IQD13, IQD31* in apple). ILR1 hydrolase regulates the rate of amido-IAA hydrolysis which results in activation of auxin signaling [42]. *ILR1* transcripts are induced by JA, suggesting that these genes might play roles in JA conjugate hydrolysis or that indoleacetic acid (IAA) release may be JA inducible. Qi et al. [43] supported the hypothesis that JA and auxin interact positively in regulating plant resistance to necrotrophic pathogens and that activation of auxin signaling by JA may contribute to plant resistance to necrotrophic pathogens. They also showed that application of IAA and methyl jasmonate (MeJA) together synergistically induces the expression of defense marker genes PDF1.2 (PLANT DEFENSIN 1.2) and HEL/PR4 (HEVEIN-LIKE), suggesting that enhancement of JA-dependent defense signaling may be part of the auxin-mediated defense mechanism involved in resistance to necrotrophic pathogens. *PR4* was found upregulated in apple. JAR1 (JA amino acid synthetase), also upregulated in apple, constitutes another important link between auxin and JA signaling [44]. Thus in apple, the JA pathway component interacting positively with auxin signaling to promote resistance against *V. inaequalis* seem to be activated.

On the contrary the remaining JA signaling (Fig. 6) seems rather repressed in apple. Indeed, numerous apple DEGs were found downregulated (*LOX1, LOX2, OPCL1, OPR2*) indicating that the JA biosynthesis was repressed in apple. However JAR1 was up regulated, probably in link with the auxin signaling as just seen before. The endogenous bioactive form of jasmonate is not JA itself but (+)-7-iso-Jasmonoyl-L-isoleucine (JA-Ile), which is produced from JA by the enzyme JAR1 [45]. In pear, *LOX5* and *ACX1* were down regulated but *OPR2* and *JMT* were up regulated. JMT encodes a JA carboxyl methyl transferase which converts JA into MeJA. Seo et al. [46] proposed that JMT is a key enzyme for the jasmonate-regulated plant responses and that MeJA can act as an intra-cellular regulator, a diffusible intercellular signal transducer, or an airborne signal mediating intra- and interplant communications. Some signals generated during defense responses may activate JMT that can self-amplify, stimulate, or regulate its own expression, propagating the MeJA-mediated cellular responses throughout whole plants. The signaling pathway depending on JA was also contrasted between apple and pear (Fig. 6). MYC2, as well as its close homologs MYC3 and MYC4, are central positive transcriptional regulators of jasmonate signaling and are negatively regulated by JAZ proteins through protein interactions. *JAZ1* was found up-regulated in apple, and thus was a negative regulator of *at4g17880* (MYC4), which was indeed down-regulated. Unlike in apple, a JAZ coding gene, *JAZ3*, was down-regulated in pear. *MYC4* was also found repressed in pear. Nevertheless, UBP12, which is known as a stabilizer of another MYC protein: MYC2 [47], was found down regulated in apple and up regulated in pear. *UBP12* down regulation reinforces the inactivation of MYC4 in apple. The inhibition of *JAZ3* and the induction of *UBP12* are consistent with the positive regulation of JA in pear. Moreover WRKY70, which is an inhibitor of the JA defense pathway, was found down-regulated in pear. Consistently with the previous hypothesis of a repressed JA signaling pathway in apple, *WRKY70* was up-regulated in that species. Among JA-responsive genes, the pathogenesis-related PR3, PR4, PR12 act downstream MYC2 activation [48]. In our data, in accordance with the repression of the inhibitor WRKY70, the *PR3* gene was found up-regulated in pear. PR3 encodes a basic chitinase involved in ethylene/jasmonic acid mediated signaling pathway during SAR. PRB1 encodes a basic PR1-like protein and is responsive to ethylene and MeJA [49]. PRB1 was also found induced in pear. To conclude, unlike in apple where only the JA pathway component interacting positively with auxin signaling seems to be activated, the global JA signaling pathway appeared rather induced in pear.

**Fig. 6:**
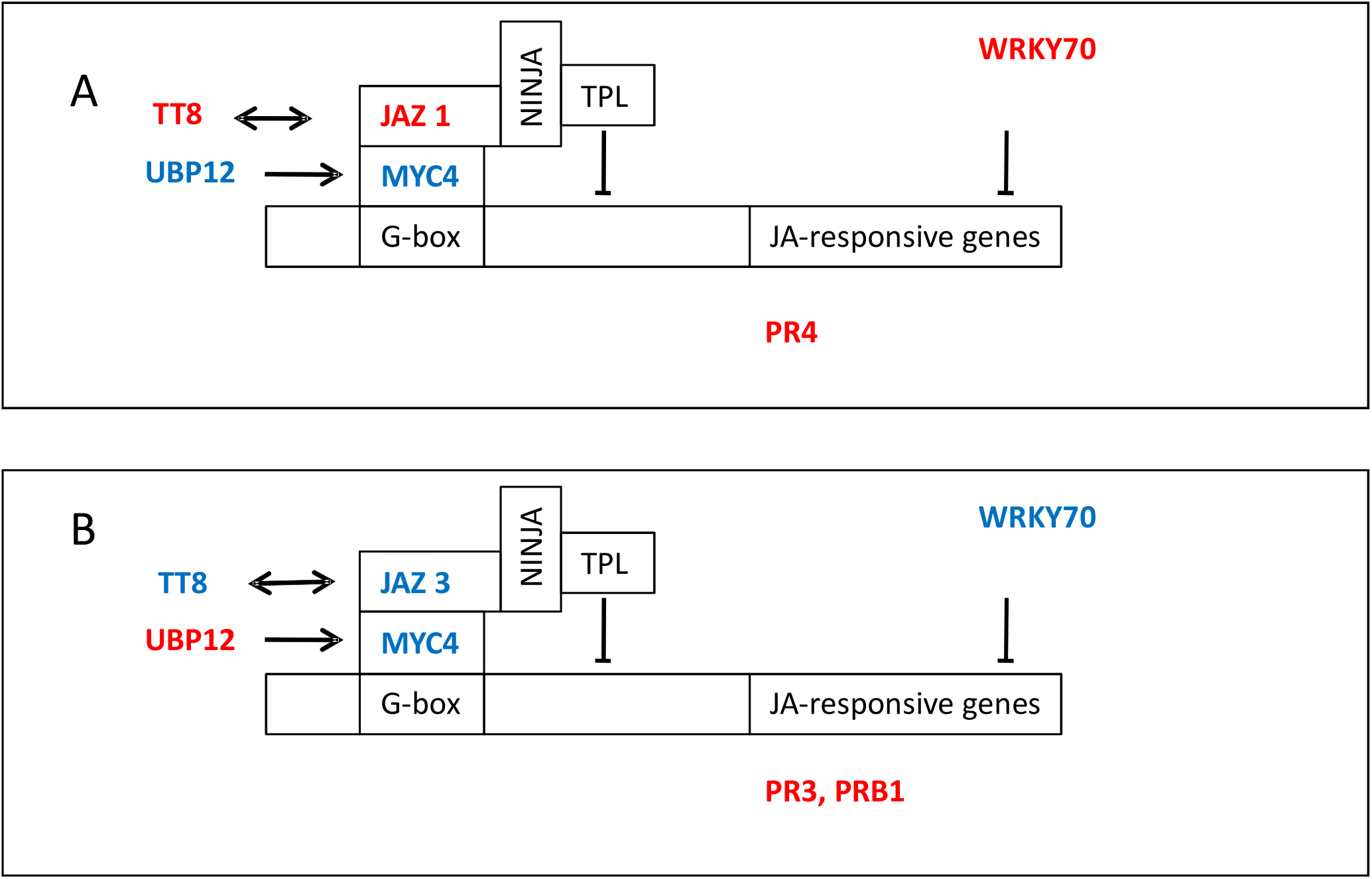
Model of JA signaling during host resistance of apple and pear to *Venturia* sp. Apple (A) and pear (B). In red: Up-regulated DEGs; in blue: down-regulated DEGs. Abbreviations : G-box: cis-element in the promoter; JAZ: jasmonate-zim domain protein; JMT: jasmonic acid carboxyl methyltransferase; MYC2: transcription factor; NINJA: novel interactor of JAZ; PR3, PR4 (HEL), PRB1 pathogenesis-related proteins; UBP12: ubiquitin-specific protease 12; TPL: TOPLESS co-repressor; TT8: Transparent Testa 8; WRKY: transcription factor.

In the BR pathway, most of the DEG found were up-regulated in apple and pear. These genes were involved in biosynthesis (*STE1, SQE1* and *CYP90A*), in signaling (*SERK1, NIK1, BAK1* in apple; *BIR3, HERK1* in pear) and regulation (*BIM2, BZR1, BEH4* in apple; *BIM2* in pear) of BRs. Anwar et al. [50] assessed that BR enhance plant tolerance to biotic and abiotic stresses, through a complex pathway to regulate the plant defense system, by activating BZR1/BES1 transcription factors. BIM2 (BES1-interacting Myc-like protein) was upregulated in apple and pear and BZR1 and BEH4 (BES1/BZR1 Homolog 4) were upregulated in apple. BR play an essential role in biotic stress tolerance by activating enzymes, biotic resistance genes, antioxidants, hormones, transcriptional factors, and signaling pathways to reduce biotic stress damage. However, BR induced resistance does not require SA biosynthesis and activation of PR gene expression, indicating that BR mediated resistance is independent of SA mediated defense signaling in plants [51]. So, some DEGS seem to indicate that BR signaling is implicated in apple and pear scab resistance.

Concerning the SA pathway, the DEGs observed were mostly negative regulators of SA (FAR1, SSI2/FAB2, CSB3/HBS, VTC2 and 5 in pear and FAR1, SSI2/FAB2, ILR3, PAP2 and VTC2 in apple) and were all activated. Wang et al. [52] demonstrated that FHY3 (Far-Red Elongated Hypocotyl 3) and FAR1 (Far-Red Impaired Response 1) negatively regulate SA signaling and plant immunity by regulating HEMB1 (which encodes a 5-aminolevulinic acid dehydratase ALAD) expression, thus providing a possible molecular linkage between light signaling and plant immunity. Gil et al. [53] showed that enhanced resistance of *csb3* (*constitutive subtilisin3*) mutant Arabidopsis plants to biotrophic pathogens requires intact salicylic acid (SA) synthesis and perception. The VTC2 and 5 genes have GDP-L-Gal phosphorylase activity and are required for L-Ascorbic Acid synthesis. MeJA treatment increased expression of VTC1 and VTC2 transcripts and enhanced L-ascorbic acid accumulation in Arabidopsis [54]. Moreover Mukherjee et al. [55] suggested that *vtc1-1, vtc2-1*, and *vtc3-1* mutants can induce defense against *Pseudomonas syringae* more rapidly through SA-dependent upregulation of pathogenesis related (PR) genes and that resistance of *vtc1* and *vtc2* mutants was due to elevated SA. SSI2/FAB2 encodes stearoyl–acyl carrier protein desaturase (S-ACP-DES). The S-ACP-DES–derived fatty acid (FA) 18:1 or its derivative is required for the activation of certain JA-mediated responses and the repression of the SA signaling pathway. Kachroo et al. [56] clearly demonstrated the importance of fatty acids in modulating signaling between SA- and JA-dependent defense pathways. These numerous negative regulators of the SA accumulation/pathway in apple and pear suggest that, even if SAR seems to be engaged (detailed later), it must be independently of SA pathway and in favor of the JA pathway.

The SA and JA/ethylene (ET) pathways are known as mutually inhibitory in many cases and important for immunity against necrotrophic and biotrophic pathogens, respectively [57]. However, Tsuda et al. [58] showed that, as both the SA and JA/ET pathways positively contribute to immunity, the loss of signaling flow through the SA pathway can be compensated by a rerouting signal through the JA/ET pathways. These findings by Tsuda et al. could explain the absence of SA pathway and the activation of the JA pathway observed in pear *Rvi6* resistance against the hemi-biotrophic fungus *V. pyrina*. BR signaling seems also implicated in that resistance. Regarding apple *Rvi6* resistance against the hemi-biotrophic fungus *V. inaequalis*, defenses seem surprisingly rather deployed independently of the signaling pathways based on the main defense hormones JA and SA, except a JA pathway component interacting positively with auxin signaling, and possibly also thanks to BR signaling.

#### Major role of cell wall-related gene modulation in both species

The plant cell wall is the first contact point during biotic stress and plays an important role in the activation and regulation of defense response strategies. The primary cell wall consists mainly of carbohydrate-based polymers (cellulose, pectins and hemicelluloses). The secondary cell wall also contains cellulose, but is enriched in lignin and xylans. The main DEGs detected at 8, 24 or 72 hpi during apple (GalaRvi6 / Gala) and pear (60AU / Conference) responses to *V. pirina* and *V. inaequalis*, respectively are listed in Table S3.

Remodeling of primary and secondary cell walls by impairing the function of cellulose synthase (CESA) genes has a specific impact on pathogen resistance and tolerance to abiotic stresses. Arabidopsis CESA mutants, like CESA3-defective isoxaben resistant (*ixr1*), illustrate that specific cell wall damage (CWD) of either the primary or secondary cell wall causes differential wall alterations, activating distinct immune responses [29]. In our study, *CESA3* was up-regulated in pear, *CESA9* was up-regulated in pear and apple, whereas *CESA6/IRX2* was down-regulated in apple. Arabidopsis GAE1 and GAE6 are required for pectin biosynthesis, leaf flexibility, and immunity to *Pseudomonas syringae* pv *maculicola* ES4326 and to specific *Botrytis cinerea* isolates [59]. These genes were up-regulated in apple and pear respectively and thus maybe involved in scab resistance. Pectin methylesterase (PME) control the esterification status of pectin. In our study, *ATPME3* was up-regulated in pear, *PMEPCRF* was down-regulated in pear, and *PME31* was up-regulated in apple. The determinant of immunity to *Alternaria brassicicola* and *Pseudomonas syringae* pv. *maculicola* ES4326 is responsible for changes in the pattern of pectin methylesterification [60]. However, the complexity of pectins is such that we are still far from understanding the details of their contribution to cell-wall integrity mediated immunity [29]. Hemicelluloses are plant cell wall polysaccharides that have β-1,4-linked backbones with an equatorial configuration [29]. Polysaccharides such as xylan, xyloglucan, mannans, glucomannans and β-1,3-1,4-glucans can be grouped under this definition [61]. L-Fucose is involved in the biosynthesis of several cell wall polymers (e.g. pectin, xyloglucan). Zhang et al. [62] found that a defect in L-fucose synthesis has a broad effect on multiple aspects of plant defense, including stomatal defense, apoplastic defense, PTI and ETI. Further analysis of known alleles of SCORD6, also named MUR1, as well as mutants of specific fucosyltransferases showed a requirement of intact N-glycans, O-glycans and mono-O-fucosylated proteins for proper *Arabidopsis thaliana* immune responses. In our study, *MUR1*, *MUR2* and *MUR3* were up-regulated in pear and *MUR2* was up-regulated in apple. The alteration of wall xylose content, the moiety that is present in xylans and xyloglucans, also affects the resistance of *Arabidopsis* to pathogens. For example, plants with enhanced levels of wall-bound xylose, as occurs in the *Arabidopsis irx6* mutants showed an enhanced resistance to the necrotrophic fungus *Plectosphaerella cucumerina* [40]. In our study, *IRX6/COBL4* was down-regulated in pear and could be involved in scab resistance. No callose synthase genes were differentially expressed in our data. The genes related to lignin will be discussed later.

Our overall results indicate that cell wall genes involved in pectin, cellulose and hemicellulose synthesis and polysaccharide degradation were mostly up-regulated during Rvi6-mediated apple and pear scab resistance, in agreement with many reports on the involvement of most of these genes in the response of plants to pathogens.

#### Importance of lipid metabolism for cuticle biosynthesis and SAR signaling

A large number of genes involved in lipid metabolism were up-regulated in pear (39 out of 50 DEGs). Most of these up-regulated DEGs were involved in fatty acid (FA) synthesis, like *ACC1* (acetyl-CoA carboxylase 1), *SSI2* (stearoyl-[acyl-carrier-protein] desaturase), *KAS1* (β-ketoacyl-ACP synthase 1), *KCS2* (3-ketoacyl-CoA synthase), *LACS2* (long-chain acyl-coenzyme A synthetase 2). Similarly, many genes involved in lipid degradation were also up-regulated (10 out of 15 DEGs), like *CER4* (alcohol-forming fatty acyl-CoA reductase). One lipid transfer protein (*LTP1*) was also up-regulated. On the contrary in apple, three genes involved in glycolipid synthesis (out of three DEGs) were up-regulated, but none of them in pear. Two of these genes were *MGD1* (monogalactosyl diacylglycerol syntase 1) and *DGD2* (digalactosyldiacylglycerol synthase 2).

A large number of studies have revealed the role of lipids and lipid metabolites during plant-pathogen interactions: (i) through the lipoxygenase pathway with the production of oxylipins and for example JA, which are important signaling molecules for defense regulation (ii) through the unsaturated FA pathway by the remodeling of membrane lipid composition and defense signaling, and finally (iii) through the very long chain fatty acid (VLCFA) pathway. The VLCFAs are FAs containing 20 to 36 carbons synthesized in the endoplasmic reticulum, which are crucial for a wide range of biological processes in plants. These lipids are indeed required for the biosynthesis of the plant cuticle [63]. Our results indicate that in pear many genes involved in the unsaturated FA and VLCFA pathways were up-regulated (Fig. 7 A). The very long CoA chains are subsequently converted to aliphatic esters, primary alcohols, alkanes or aldehydes by one of the two wax biosynthetic pathways. CER4 encodes FA-CoA reductase and a mutation in *cer4* leads to a major decrease in primary alcohols and wax esters. A second branch of wax biosynthetic pathway is responsible for the formation of alkanes, secondary alcohols and ketones. CER1 likely encodes a decarbonylase and is involved in alkane formation. In comparison to cuticular wax biosynthesis, monomers of cutin are synthesized from C16:0 and C18:1 FAs. LACS2 encoded acyl-CoA synthetase may either be required to synthesize 16-hydroxy 16:0-CoA, a substrate for ω-hydroxylase, or required for membrane transfer of cutin monomers [64]. Throughout the FAs synthesis pathway, several genes were up-regulated in pear leading to synthesis of cuticular waxes and cutin, components of cuticle (Fig. 6A). Cutin and cuticular waxes are known to serve as a physical barrier against pathogens.

**Fig. 7:**
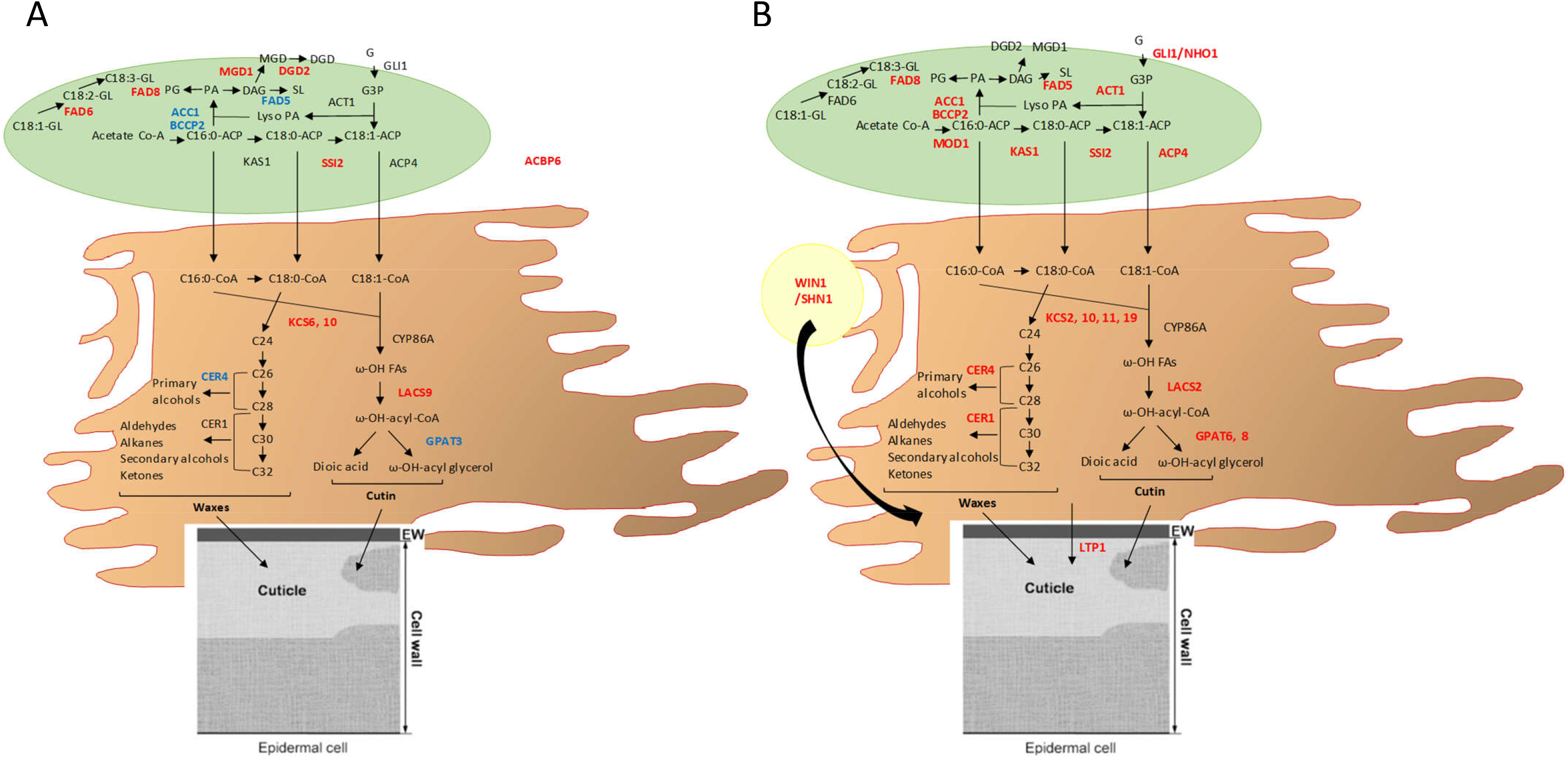
Model of lipid metabolism and cuticle biosynthesis during host resistance of apple and pear. (adapted from [64], [118] and [119]). Apple (A) and pear (B). In red: up-regulated DEGs; in blue: down-regulated DEGs. Abbreviations: ACBP, acyl CoA binding protein; ACC1, acetyl-CoA carboxylase 1; ACP, acyl carrier protein; BCCP2, biotin carboxyl carrier protein 2; CER, alcohol-forming fatty acyl-CoA reductase; CoA, coenzyme A; DAG, diacylglycerol; DGDG, digalactosyldiacylglycerol; FAD, fatty acid desaturase; GLI1, glycerol kinase; GPAT, glycerol-3-phosphate acyltransferase; KAS, β-ketoacyl-ACP synthase; KCS, 3-ketoacyl-CoA synthase; LACS, long-chain acyl-coenzyme A synthetase; LTP1, lipid transfer protein 1; MGDG, monogalactosyldiacylglycerol; MOD1, mosaic death 1; PA, phosphatidic acid; PG, phosphatidylglycerol; SL, sulfolipid; SSI2, stearoyl-ACP desaturase; WIN1/SHN1, wax inducer 1/shine 1.

Several transcription factors (TFs) have been shown to regulate cuticle biosynthesis. The most studied is SHINE1/WAX INDUCER1 (SHN1/WIN1) which is a member of the plant-specific family of AP2/EREBP transcription factors [65]. In our study, *SHN1/WIN1* TF was up-regulated in pear. Non specific Lipid Transfer Proteins (nsLTPs) play a key role in plant resistance to biotic and abiotic stresses. This justifies the classification of nsLTPs among the pathogenesis-related proteins. Indeed, LTP1 is predicted to be a member of PR-14 pathogenesis-related protein family. In our study, *LTP1* was up-regulated in pear. Two main basic mechanisms are probably involved, which agrees with the extracellular location of nsLTPs: (1) the formation of hydrophobic protective layers (cutin and suberin); and (2) the inhibition of fungal growth. Based on the binding properties of FA derivatives, a role has been proposed for nsLTPs in cutin-monomer transport during cuticle formation. Although the precise role of nsLTPs in the formation of cutin layers is unknown, they could play a role in transporting hydrophobic monomers to the extracellular polymerization hydrophobic–hydrophilic cutin–cell-wall interface where they are usually found [66].

Concerning the FAs and VLCFAs pathways, our results indicate that most of these genes were down-regulated in apple (Fig. 7 B). However, unlike pear *FAD6, MGD1* and *DGD2* were up-regulated. The FAD6 and FAD7/FAD8 enzymes can act on glycerolipids containing either C16 or C18 FAs. Enzymes catalyzing galactolipid biosynthesis are present in the inner [monogalactosyl synthase (MGD)] and outer [digalactosyl synthase (DGD)] membranes of the chloroplast and catalyze the biosynthesis of monogalactosyldiacylglycerol (MGDG) and digalactosyldiacylglycerol (DGDG), respectively. MGDG and DGDG are required for thylakoid formation. Chaturvedi et al. [67] showed that SAR, but not basal resistance of *Arabidopsis* to a *P. syringae* pv. *maculicola*, was compromised in the *mgd1* mutant. SAR is an inducible defense mechanism that confers resistance against a broad spectrum of pathogens. They suggested that a plastid synthesized glycerolipid, most likely a galactolipid, is required for the establishment of SAR.

SAR signal generation and perception requires an intact cuticle, a waxy layer covering all aerial parts of the plant [68]. Xia et al. [64] showed that mutations in ACP4, a critical component of FA biosynthesis, impairs systemic acquired resistance (SAR) because it affects cuticle formation in the leaf. SAR is also compromised in *lacs2, lacs9, cer1, cer3* and *cer4* mutants. These results suggest that perception of the mobile signal by the cuticle in distal leaves is as important as its generation at the site of the primary infection, as an intact cuticle is required for the perception of the mobile SAR signal. They also showed that the acyl CoA binding protein ACBP6, a soluble protein that is present in the cytosol, may be involved in the transport of FAs and/or lipid species required for the proper development of the plant cuticle as well as generation of the mobile SAR signal [69]. They suggested that *acbp* plants are defective in the generation of SAR signal but competent in its perception [64]. *ACBP6* gene was up-regulated in apple, consequently *Rvi6* apple lines could generate the mobile SAR signal. Whereas *ACP4* was up-regulated in pear. This suggest that *Rvi6* pear could be competent in the perception of the mobile SAR signal by the cuticle in distal leaves. SAR is also positively regulated by CRY1, upregulated in apple and pear. Thus, despite the fact that the SA pathway was not activated, apple and pear seem to engage SAR. Moreover Kachroo and Robin [70] showed that SA is well known to accumulate in the primary infected leaf, but there is little to no evidence supporting that SA accumulation in the distal tissues is required for the induction of SAR. The connection with the JA could be made as well in pear. Indeed JA has been suggested to participate in SAR [71]. *PR3* gene was found up-regulated in pear and encodes a basic chitinase involved in ET/JA mediated signaling pathway during SAR. Moreover exogeneous application of MeJA is known to induce SAR [72]. In pear, via the induction of JMT, MeJA could be produced.

Our overall results indicate the importance of lipid metabolism for the two species. In pear, the biosynthesis of cutin and cuticular waxes is increased, which strengthen this barrier against pathogens. In apple, galactolipid biosynthesis seems to be required for the establishment of SAR and thus for resistance to scab. Furthermore, SAR signaling is activated in both species. Whereas the generation of the mobile signal is increased in apple, the perception of this signal in distal tissues is increased in pear, maybe via JA signaling.

#### General activation of the phenylpropanoid pathway and specific metabolite production in apple and pear

The phenylpropanoid pathway is involved in lignin, flavonoid and other metabolites biosynthesis. In our study, genes involved in these two first branches of the phenylpropanoid pathway were up-regulated in pear, 9 out of 9 DEGs and 8 out of 9 DEGs respectively. Most of these genes were expressed as early as 8 hours post inoculation. The trend was more towards down-regulation in apple. Lignin reinforces the cell wall, facilitates water transport and acts as a physical barrier to pathogens. Flavonoids can also have important functions as defense compounds. Indeed, flavonoids are a group of polyphenol compounds with known antioxidant activities. Flavonoids constitute a diverse family that can be divided into sub-families including flavonols, flavanols, flavanones, anthocyanidins, chalcones and dihdrochalcones. Dihydrochalcones are mainly found in apple leaves (*Malus domestica*) [73].

In our results, among the apple DEGs appeared a number of regulators of the phenylpropanoid pathway. This pathway is regulated by MYB transcription factors and Ma and Constabel [74] described some of the MYB repressors implicated. *At*MYB4 is a MYB repressor of the C4H gene of the lignin pathway. In our study, this gene was down-regulated in apple, therefore it is not repressing the lignin pathway. Indeed, the C4H gene was up-regulated in apple and also in pear. *At*CPC is a MYB repressor of the anthocyanin pathway [75]. This gene was up-regulated in apple, which suggests that the anthocyanin pathway was therefore not activated (Fig. 8 A). Moreover Qi et al. [76] revealed that JAZ proteins interact with bHLH (Transparent Testa8, Glabra3 [GL3], and Enhancer of Glabra3 [EGL3]) and R2R3 MYB transcription factors (MYB75 and Glabra1), essential components of WD-repeat/bHLH/MYB transcriptional complexes, to repress JA-regulated anthocyanin accumulation and trichome initiation. In our study *TT8* and *JAZ1* were down regulated in apple. Thus, we can assume that the anthocyanin pathway was not activated in apple scab resistance. Concerning the flavonoid pathway, *CHS* and *GT72B1* were up-regulated in apple. This suggest that phlorizin could be synthetized from phloretin by glycosylation of GT72B1 and involved in the resistance to apple scab. Indeed, Gaucher et al. [73] showed the potential role of dihydrochalcones in protecting apple tissues against biotic stresses, in particular *Erwinia amylovora* infection, another major pathogen of apple trees. Judgé et al. [77] identified a novel uridine 5’-diphospho-glucuronyl transferase (UGT) from apple, *MdPGT1*, belonging to the UGT88 family, using a functional genomic approach. Silencing of *UGT88F1*, a phloretin-specific glycolsyltransferase, perturbs the phenylpropanoid pathway. Metabolite analysis of the UGT88F1 knockdown lines revealed a number of quercetin, kaempferol and phloridzin glycosides that were significantly reduced in leaves [78]. Even if *MdPGT1* clusters with members of the UGT88 family and separately from other Arabidopsis UGT sequences, *GT72B1* was clearly over-expressed in apple in our study and could be involved in phlorizin biosynthesis.

**Fig. 8:**
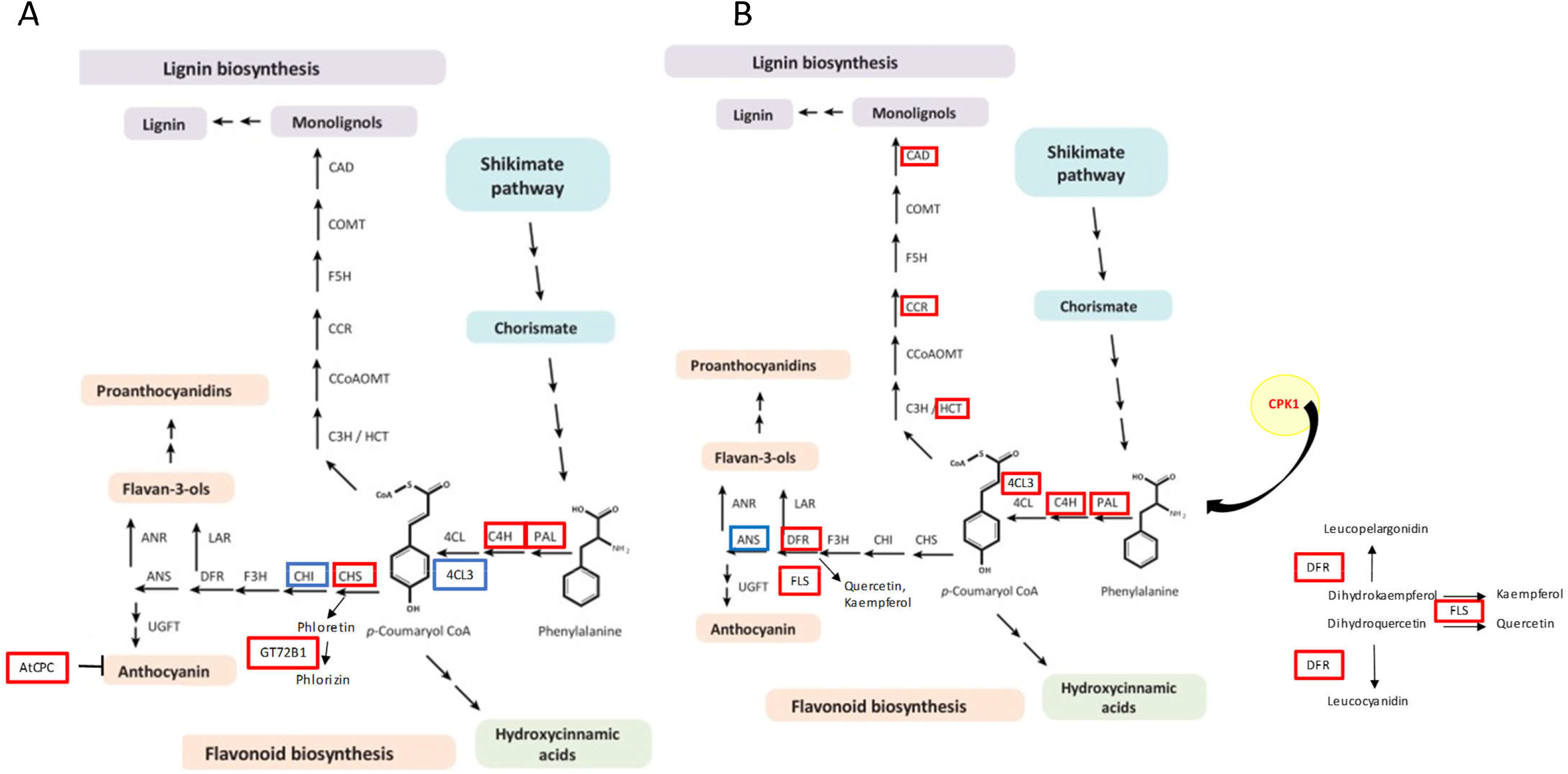
Overview of the phenylpropanoid pathway during host resistance of apple and pear to *Venturia* sp. (adapted from [74]). Apple (A) and pear (B). In red: up-regulated DEGs; in blue: down-regulated DEGs. Abbreviations: 4CL, 4-coumarate-CoA ligase; ANR, anthocyanidin reductase; ANS, anthocyanin synthase; C3H, cinnamate 3-hydroxylase; C4H, cinnamate 4-hydroxylase; CAD, cinnamyl alcohol dehydrogenase; CCoAOMT, caffeoyl-CoA O-methyltransferase; CCR, cinnamoyl-CoA reductase; CHI, chalcone isomerase; CHS, chalcone synthase; COMT, caffeic acid 3-O-methyltransferase; CPK, calcium-dependent protein kinase; DFR, dihydroflavonol reductase; DIR, plant dirigent proteins; F3H, flavanone 3-hydroxylase; FLS, flavonol synthase; GT72B1, UDP-glycosyltransferase; HCT, hydroxycinnamoyl-CoA shikimate/quinate hydroxycinnamoyl transferase; KFB, Kelch repeat F-box; LAR, leucoanthocyanidin reductase; OMT1, O-methyltransferase 1; PAL, phenylalanine ammonia-lyase; UGFT, UDP-glucose flavonoid-3-O-glucosyltransferase; UGT72E1, coniferyl-alcohol glucosyltransferase.

In pear, the lignin pathway was activated with many genes up-regulated all along this pathway. Phenylalanine ammonia-lyase (PAL) is the key enzyme of phenylpropanoid biosynthesis and is subject to post-translational phosphorylation. The phosphorylation of PAL is accomplished by the activation of *AtCPK1* [79], which was up-regulated in pear in our experiment (Fig. 8 B). The *C4H* and *4CL3* genes were also up-regulated, leading to either flavonoid or lignin pathways. Several enzymes are encoded by multigene families, in which the different members may have different specific activities and tissue/cell-specific expression patterns. For instance, Arabidopsis 4CL1 and 4CL2 are primarily involved in the biosynthesis of p-coumaroyl-CoA for lignin, and 4CL3 in that of p-coumaroyl-CoA for flavonoids [80]. Even if the flavonoid pathway could be favored, the lignin biosynthesis seems also to be engaged with many genes up-regulated all along this pathway, in particular *CAD7* which is involved in the synthesis of the lignin precursor. Lignin is an aromatic polymer that is mainly deposited in secondary thickened cell walls. Increased accumulation of lignin can provide a basic barrier against pathogen spread and reduce the infiltration of fungal enzymes and toxin into plant cell walls [81].

The importance of the regulation by the SCF E3 ubiquitin ligase complex in apple and pear was already presented before. Among the E3-SCF genes, 2 Kelch repeat-containing F-box (KFB) family protein were up-regulated in pear, one of which was highly up-regulated (ratio of 3.16). The Kelch motif containing F-box (KFB) proteins are a subfamily of F-box proteins. KFB^CHS^ (chalcone synthase) and KFB^PAL^ are clustered in a same clade sharing 31 to 36% overall sequence similarity. However, KFB^CHS^ has little propensity to interact with other phenylpropanoid pathway enzymes excepting for chalcone synthase (CHS) [82]. The gene *at1g23390* overexpressed in pear (named *KFB*) physically interacts with *CHS* and specifically mediates its ubiquitination and degradation [82]. That could explain why no *CHS* was activated in pear.

However in pear, the flavonoid pathway was also activated, with the up-regulation of dihydroflavonol 4-reductase (DFR) and flavonol synthase (FLS) genes. The FLS gene leads to the biosynthesis of quercetin and kaempferol. These metabolites are activated after priming of tomato seeds with meJA in response to *Fusarium*, in rust infected leaves of black poplar, are involved in pecan scab resistance, and in Norway spruce rust resistance [83, 84, 85, 86]. Therefore, flavonoids, more specifically quercetin and kaempferol, could be involved in pear scab resistance.

Our overall results indicate that most of the flavonoid and lignin genes were up-regulated in pear while as many genes were up-regulated as down-regulated in apple. Lignin is an important barrier against pests and pathogens. A general positive correlation between lignin amount and pathogen resistance is observed for several plant-pathogen interactions (*Fusarium* sp., *Xanthomonas* sp., *Verticillium* sp., *Pseudomonas syringae* pv. *Tomato*) [87]. Flavonoids are also involved in resistance, as previously reported in apple. Indeed, accumulation of phloridzin and phloretin proved to be related with quantitative resistance to *V. inaequalis*, suggesting the creation of a harmful environment [88, 89]. Gaucher et al. [73] showed that dihydrochalcones accumulate mainly in the upper layers of apple leaves as well as in parenchymal non-conductive cells found in the vascular bundles. As the localization results suggest, dihydrochalcones could constitute a physico-chemical barrier against pathogens colonizing leaf upper layers, such as *V. inaequalis*.

## Conclusion

To conclude, our study allowed elucidating the mechanisms underlying the major gene *Rvi6*-induced resistance in apple, but also in pear. In apple/*V. inaequalis* interaction, once achieved the pathogen recognition thanks to *Rvi6*, signal transduction is triggered by calcium and hormonal signaling, in particular auxin and BRs. This leads to the induction of defense responses such as a slight remodeling of primary and secondary cell wall, galactolipids biosynthesis, the establishment of a SAR and the biosynthesis of flavonoids (Fig. 9). In pear/*V. pyrina* interaction, once achieved the pathogen recognition, signal transduction is triggered by calcium, ubiquitin, G protein and hormonal signaling involving brassinosteroids but especially JA. The perception of a SAR signal in distal tissues is also observed. This leads to the induction of defense responses such as the remodeling of primary and secondary cell wall, the biosynthesis of cutin and cuticular waxes and the biosynthesis of flavonoids and lignins (Fig. 9).

**Fig. 9:**
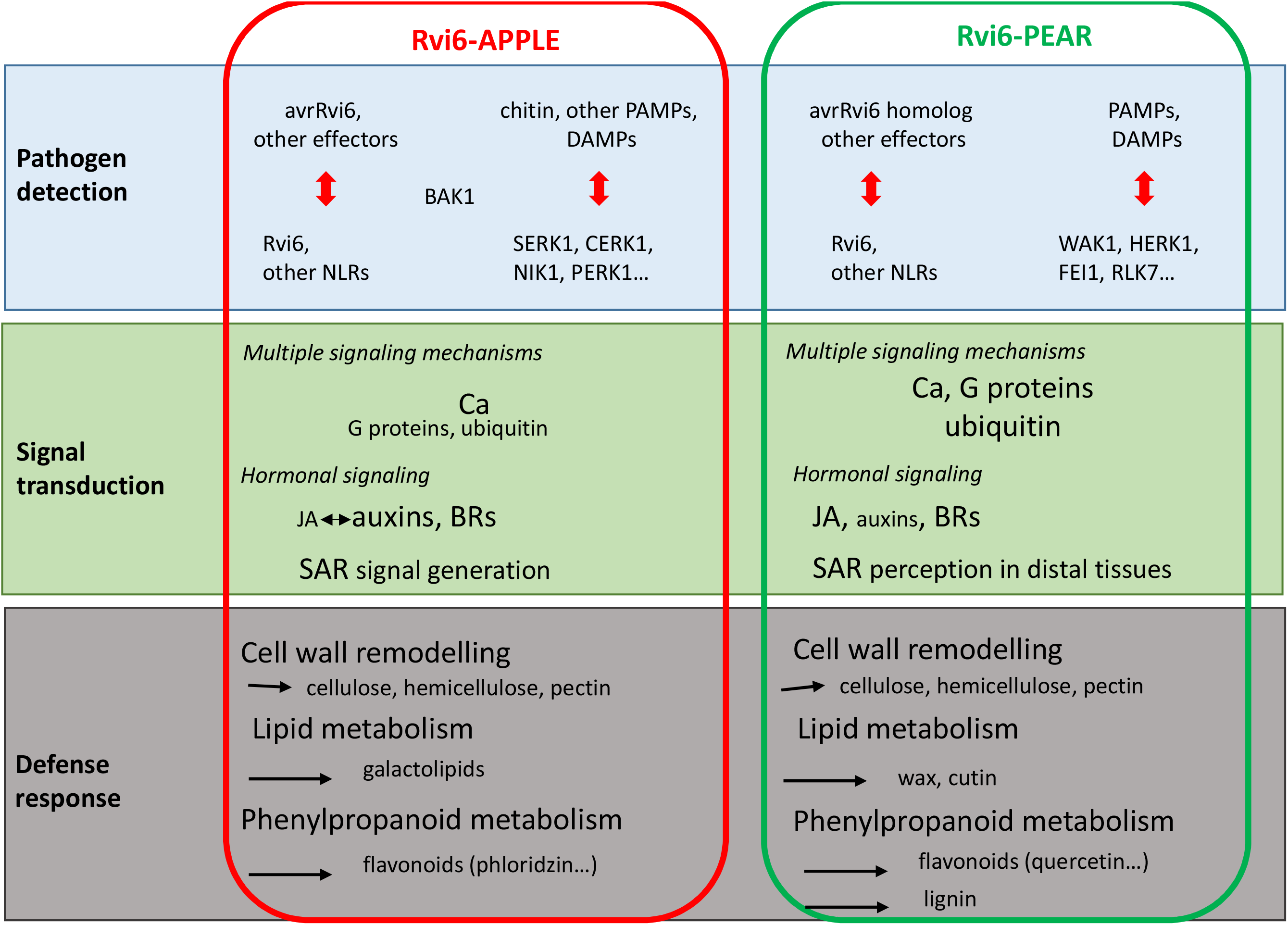
Hypothetical scheme of the main features of *Rvi6*-mediated scab resistance in apple and pear. Based on the most relevant DEGs described in the present study.

Beyond the precise deciphering of *Rvi6*-induced signaling and defense cascades in apple and pear, this work also revealed that these downstream mechanisms differ between both genera. We can venture that it is also the case at the inter- and intraspecific levels. Comparative transcriptomic analyses of *Rvi6* introgressions in various apple and pear genetic backgrounds would be a great help to test that hypothesis, which has important implications in terms of ideotype design for improved and sustainable resistance.

The present work is the first example of a successful intergeneric transfer of an NLR gene in the *Rosaceae* family. Indeed, apple and pear belong to the *Maloideae* subfamily within the *Rosaceae*, and originate from a common ancestor that probably was susceptible to *Venturia*, but they diverged about 8-16 million years ago during evolution [90], and are now hardly crossable anymore. The pathogen most probably followed both directions, adapting to apple (*V. inaequalis*) or to pear (*V. pyrina* and *V. nashicola*). *V. inaequalis* is not pathogenic on pear and *V. pyrina* not on apple [91]. This reflects the sexual incompatibility (or poor compatibility) of the hosts. One could name this non-host resistance. The present study showed both the efficiency of this transfer strategy in terms of resistance, its limits given the partial breakdown observed with the *V. pyrina* VP 98 strain, and raises questions. Indeed, *V. pyrina* has very likely not encountered *Rvi6* in apple but surprisingly, *Rvi6* provides resistance to *V. pyrina*. Even more surprising is that *V. pyrina* has a strain that has (partially) overcome the *Rvi6* resistance, although there was no expected selection on *V. pyrina* for virulence to *Rvi6*. This is very remarkable because as far as we know, pear does not have a functional *Rvi6* homolog, providing resistance to pear scab. Perchepied et al. [92] mapped scab resistance on chromosome 01 of pear (called *qrpv_LG01*), but the QTL peak is approximately 40 cM distant from the *Rvi6* syntenic region, thus in a very distinct genomic region. Apparently, the *V. pyrina* strain developed already virulence, in spite of the fact that virulence was not needed (yet). Is this virulence inherited from an ancient *Venturia* strain that had overcome the *Rvi6* resistance already before *V. inaequalis* and *V. pyrina* diverged? That explanation is only plausible if the *Rvi6* virulence does not have a negative effect on aggressiveness. Otherwise, the unneeded virulence would have been lost already during the numerous generations of the pathogen after the speciation. An alternative explanation is that pear still harbors a functional *Rvi6* homologue, but that homologue may have been still unnoticed. Another explanation could be that *V. pyrina* is continuously modifying its effectors, some of them, as for *V. inaequalis* [5, 93], are able to overcome *Rvi6*, and that we co-incidentally tested a strain with that type of effector. A strategy to achieve sustainable resistance could be to associate the *Rvi6* qualitative resistance (based on a single major gene) with quantitative resistances (based on multiple genes of partial effect, QTLs). Lasserre-Zuber et al. [94] evaluated the utility of pyramiding the *Rvi6* gene with QTLs F11 and F17, in two unsprayed orchards where resistance effectiveness of these QTLs and this major gene has already been challenged. Their results showed that the effectiveness of pyramiding to control scab was dependent on the site and could not be completely explained by the effectiveness level of the resistances alone. Moreover, diversifying resistance mechanisms is expected to promote conflicting selection pressures on the pathogen populations, which should constrain them to an evolutionary compromise limiting their development. Indeed, Laloi et al. [95] showed that the combination of QTLs acting on different stages of pathogen development, and thus probably based on different mechanisms, could improve the efficiency of resistance in the apple scab pathosystem.

Thus, the deciphering of the molecular mechanisms underlying varied resistances, quantitative as qualitative, must be pursued, in order to allow the design of efficient and sustainable host type resistances in apple and pear, allying adapted genetic background choice, pyramiding and intergeneric transfer strategies. Moreover, whatever the genetic combination used, the adaptation of the pathogen will have to be considered as well.

## Methods

### Biological material

The apple cultivar ‘Gala’ was chosen because of the availability of the transgenic clone P_MdRbc_HcrfVf2-11 (GalaRvi6) already described by Joshi et al. [10]. This transgenic clone expresses the *Rvi6* transgene under the control of the apple *Rubisco* gene promoter and terminator sequences. For this study, it was micropropagated on Murashige and Skoog [96] medium supplemented with 0.5 mg/l 6-benzyladenine and 0.1 mg/l 3-indolebutyric acid. The pear cultivar ‘Conference’ (CF) was chosen for this study because of its high susceptibility to scab [21]. This cultivar was propagated in vitro as previously reported [97].

For pear transformation, the *A. tumefaciens* strain EHA105 [98] containing the ternary plasmid pBRR1MCS-5 with a constitutive *VirG* gene [99] was used. The binary plasmid pMF1-pMdRbc1.6-Vf2-tMdRbc [10] contained the *Rvi6* coding sequence and the regulatory promoter and terminator sequences from the apple *Rubisco* gene.

For apple scab inoculation, the *V. inaequalis* monoconidial isolate used was EU-B05 from the European collection of *V. inaequalis* of the European project Durable Apple Resistance in Europe [100]. For pear scab inoculation, three monoconidial strains of *V. pyrina* were chosen for their aggressiveness on ‘Conference’ (VP98, VP102 and VP137, [21]).

### Production of pear transgenic lines and molecular characterization

Leaves from micropropagated ‘Conference’ plants were used as explants and *Agrobacterium tumefaciens*-mediated transformation was performed according to Mourgues et al. [101]. Transgenic lines and control plants were then propagated *in vitro* and acclimatized in a greenhouse as described by Faize et al. [102] for apple and Djennane et al. [103] for pear.

The ploidy level of 30 independent pear kanamycin resistant lines was checked by flow cytometry, as described in Chevreau et al. [104], and tetraploid lines were eliminated.

Presence of the transgene and absence of contaminating agrobacteria were monitored by PCR in 30 independent pear kanamycin resistant lines. Genomic DNA extraction from leaves of *in vitro* shoots was performed according to [105]. Primers used allowed the detection of (i) the specific pMdRbc1.6-Rvi6 fragment, (ii) *A. tumefaciens* presence, (iii) *nptII* gene and (iv) elongation factor 1α (*EF*1*α*). They are listed in Table S4. Amplifications were performed using GoTaq^®^ Flexi DNA Polymerase (Promega) according to the manufacturer’s recommendations. The PCR reaction conditions were identical for the four genes: 95°C for 5 min, followed by 35 cycles at 95°C for 30 s, 60°C for 45 s, 72°C for 1min 30s, with a final extension at 72°C for 5 min. The PCR products were separated on a 1.5 % agarose gel.

Expression of the transgenes was assessed by Quantitative RT-PCR (QPCR). Total RNA was extracted from leaves of greenhouse-grown plants, according to the same protocol as in 2.5 section. First-strand cDNA synthesis and QPCR were then performed as described in 2.6 section, with 0.3 (*EF1α* and pMdRbc1.6-Rvi6 fragment) of each primer (Table S4) (10 μM) in a final volume of 10 μl. Normalization was done with the reference gene *EF1α* and the non-transgenic genotype ‘Conference’ was used as a calibrator.

### Scab inoculation procedure

Greenhouse growth conditions and mode of inoculum preparation were as described in Parisi and Lespinasse [106] for apple and Chevalier et al. [107] for pear. Briefly, the youngest leaf of actively growing shoots was tagged and the plants inoculated with a conidial suspension (2 × 10^5^ conidia ml^−1^). Symptoms were recorded at 14, 21, 28, 35 and 42 days after inoculation. The type of symptoms was scored using the six class-scale of Chevalier et al. [22] and the quantity of disease was evaluated by the percentage of leaf area with sporulating lesions, recorded on a 7 class-scale as described in Parisi et al. [108].

### Microscopic observations

Two types of microscopic observations were performed. Histological studies were made on samples stained with the fluorophore solophenyl flavine [109]. In brief, leaf discs were rinsed in ethanol 50° before staining in a water solution of solophenyl flavine 7GFE 500 (SIGMA-Aldrich, St Louis USA) 0.1% (v/v) for 10 min. The samples were first rinsed in deionized water, then in glycerol 25% for 10 min. Finally, the leaf samples were mounted on glass-slides in a few drops of glycerol 50%. They were examined with a wide-field epifluorescence microscope BH2-RFC Olympus (Hamburg, D) equipped with the following filter combination: excitation filter 395 nm and emission filter 504 nm. For cryo-scanning-electronic microscopy (SEM), leaf samples were fixed in glutaraldehyde 4% in phosphate buffer 0.2M, pH 7.2 under vacuum and stored at 4°C until observation with a benchtop SEM Phenom G2 Pro (PhenomWorld, Eindhoven, NL).

### Transcriptomics experiment

Leaf samples were immediately frozen in liquid nitrogen and kept at −80°C until analysis. Sampling concerned the youngest expanded leaf of each plant labeled the day of the inoculation. Each sample is a pool of leaves from three different plants. Four biological repeats (genotype × treatment × time) were collected during two independent scab inoculation tests. Leaf samples taken just before inoculation (T0) and at 8, 24 and 72 hours post inoculation of two biological repeats among these four were then used to perform transcriptomics analyses (Table 2).

**Table 2:** Design of the transcriptomic experiments.

| Species | Genotype | Scab strain | Type of interaction | Transcriptomic analysis |
| --- | --- | --- | --- | --- |
| Pear | 60AU | VP102 | R* | R against S at T0<sup>§</sup>, 8, 24 and 72 hpi |
|  | Conference | VP102 | S** | R against S at T0, 8, 24 and 72 hpi |
| Apple | GalaRvi6 | VI EUB04 | R | R against S at T0, 8, 24 and 72 hpi |
|  | Gala | VI EUB04 | S | R against S at T0, 8, 24 and 72 hpi |

\* R: resistance, \*\* S: susceptibility, <sup>§</sup>T0: sampling time just before inoculation

For RNA extraction, frozen leaves were ground to a fine powder in a ball mill (MM301, Retsch, Hann, Germany). RNA was extracted with the kit NucleoSpin RNA Plant (Macherey Nagel, Düren, Germany) according to the manufacturer’s instructions but with a modification: 4% of PVP40 (4 g for 100 ml) were mixed to the initial lysis buffer RAP before use. Purity and concentration of the samples were assayed with a Nanodrop spectrophotometer ND-1000 (ThermoFisher Scientific, Waltham, MA, USA) and by visualization on agarose gel (1% (weight/volume) agarose, TAE 0.5×, 3% (volume/volume) Midori green). Intron-spanning primers designed on the *EF1α* gene were used to check the absence of genomic DNA contamination by PCR. The PCR reaction conditions were as follows: 95°C for 5 min, followed by 35 cycles at 95°C for 30 s, 60°C for 45 s, 72°C for 1 min, with a final extension at 72°C for 5 min. The PCR products were separated on a 2% agarose gel.

Amplifications (aRNAs) were produced with MessageAmpII aRNA Kit (Ambion Invitrogen, Waltham, MA, USA), from 300 ng total RNA. Then 5 μg of each aRNA were retrotranscribed and labelled using a SuperScript II reverse transcriptase (Transcriptase inverse SuperScript™ II kit, Invitrogen, Carlsbad, CA, USA) and fluorescent dyes: either cyanine-3 (Cy3) or cyanine-5 (Cy5) (Interchim, Montluçon, France). Labeled samples (30 pmol each, one with Cy3, the other with Cy5) were combined two by two, depending on the experimental design. For each comparison two biological replicates were analysed in dye-switch as described in Depuydt et al. [110]. Paired labeled samples were then cohybridized to Agilent microarray AryANE v2.0 (Agilent-070158_IRHS_AryANE-Venise, GPL26767 at GEO: https://www.ncbi.nlm.nih.gov/geo/) for apple, or Pyrus v1.0 (Agilent-078635_IRHS_Pyrus, GPL26768 at GEO) for pear, containing respectively 133584 (66792 sense and 66792 anti-sense probes) and 87812 (43906 sense and 43906 anti-sense probes) 60-mer oligonucleotide probes. The hybridizations were performed as described in Celton, Gaillard et al. [111] using a MS 200 microarray scanner (NimbleGen Roche, Madison, WI, USA).

For microarray analysis we designed two new chips. For apple we used a deduplicated probeset from the AryANE v1.0 ([111]; 118740 probes with 59370 in sense and 59370 in anti-sense) augmented by 14844 probes (7422 in sense and 7422 in anti-sense) designed on new gene annotations from *Malus domestica* GDDH13 v1.1 (https://iris.angers.inra.fr/gddh13 or https://www.rosaceae.org/species/malus/malus_x_domestica/genome_GDDH13_v1.1). These probes target new coding genes with UTRs when available, manually curated micro-RNA precursors and transposable elements. For transposable elements we used one consensus sequence for each family and a randomly peaked number of elements proportionally to their respective abundance in the genome. The microarray used in this study also have probes for coding genes of *V. inaequalis* but they have not been taken into account in this study.

For pear the design was done on the *Pyrus communis* Genome v1.0 Draft Assembly & Annotation available on GDR (https://www.rosaceae.org/species/pyrus/pyrus_communis/genome_v1.0) web site. We have downloaded the reference genome and gene predictions fasta files and structural annotation gff file the 21st of September 2015. Using home-made Biopython scripts we have extracted spliced CDS sequences with 60 nucleotides before start and after stop codons to get UTR-like sequences likely to be found on transcripts resulting in a fasta file containing 44491 sequences. These 60 nucleotides size increase the probability of finding specific probes on genes with high similarity. This file was sent to the eArray Agilent probe design tool (https://earray.chem.agilent.com/earray/) to generate one probe per gene prediction. Options used were: Probe Length: 60, Probe per Target: 1, Probe Orientation: Sense, Design Options: Best Probe Methodology, Design with 3’ Bias. The probeset was then reverse-complemented to generate anti-sense probes and filtered to remove duplicated probes. The final probeset contains 87812 unique probes targeting 1 (73612 probes) or more (14200 probes) potential transcript both in sense and anti-sense.

### QPCR validation of transcriptomic data

In order to validate transcriptomic data, QPCR was performed on a selection of gene/sample associations (Table S2). First-strand cDNA was synthesized using total RNA (2.0 μg) in a volume of 30 μl of 5× buffer, 0.5 μg of oligodT15 primer, 5 μl of dNTPs (2.5 mM each), and 150 units of MMLV RTase (Promega, Madison, WI, USA). The mixture was incubated at 42°C for 75 min.

QPCR was then performed. Briefly, 2.5 μl of the appropriately diluted samples were mixed with 5 μl of PerfeCTa SYBR Green SuperMix for iQ kit (Quantabio, Beverly, MA, USA) and 0.1 or 0.2 μl of each primer (10 μM) in a final volume of 10 μl. Primers were designed with Primer3Plus or by hand, their volumes were according to their optimal concentration (determined for reaction efficiency near to 100%; calculated as the slope of a standard dilution curve; [112]). Accessions, primer sequences and optimal concentrations are indicated in Table S2. The reaction was performed on a CFX Connect Real-Time System (BIO-RAD, Hercules, CA, USA) using the following program: 95°C, 5 min followed by 40 cycles comprising 95°C for 3 s, 60°C for 1 s. Melting curves were performed at the end of each run to check the absence of primer-dimers and nonspecific amplification products. Expression levels were calculated using the ΔΔCT method [113] and were corrected as recommended in Vandesompele et al. [114], with three internal reference genes (GADPH, TUA and ACTIN 7 for apple, GADPH, TUA and EF1α for pear) used for the calculation of a normalization factor. For each couple DEG/sample (sample defining a plant, time, treatment and biological repeat combination), the ratio is gotten by dividing the mean value of CT calculated from 3 technical repeats by the normalization factor obtained for this sample.

### Statistical analyses

For scab inoculation results, the AUDPC based on sporulation scores at 14, 21, 28, 35 and 42 days after inoculation. Statistical analyses were performed with XLSTAT by using the nonparametric Wilcoxon test (p < 0.05).

Normalization and statistical analyses performed to get normalized intensity values have been done as in Celton, Gaillard et al. [111]. For each comparison and each probe, we retrieved a ratio of the logarithms of the fluorescence intensities (one per compared sample, cf. Table 2) and an associated p-value. The applied p-value threshold to determine DEGs (differentially expressed genes) was 0.05. Through blast analysis, a TAIR accession number (The Arabidopsis Information Resource; https://www.arabidopsis.org/; [115]) has been linked to a majority of apple or pear “probe/corresponding gene” and the couple “TAIR accession/ratio value” has then been used to make a global analysis of functional categories observed in the Mapman software (https://mapman.gabipd.org/homemapman.gabipd.org; [116]). The detailed analysis of DEGs has been done through TAIR and KEGG (https://www.genome.jp/kegg/) databases, and bibliography. Metadata for the DEGs discussed in this work are available in Table S5 (Online only).

## Supplementary Information

**Additional File 1: Table S1:** Scab qualitative note of nine transgenic pear lines and non-transgenic Conference inoculated with three *V. pirina* strains. Percentage of plants in the different classes of symptoms, 42 days after inoculation. **Table S3:** Expression modulation of cell wall related DEGs detected at 8, 24 or 72 hours post-inoculation during apple (GalaRvi6 / Gala) and pear (60AU / Conference) responses to *V. pirina* and *V. inaequalis*, respectively. In red: up-regulated DEGs, in blue: down-regulated DEGs.

**Additional file 2: Table S2:** DEGs analysed by QPCR. **Table S4:** Accessions and primer sequences for molecular characterization of pear transgenic lines.

**Additional File 3: Fig. S1:** Functional categories of DEGs at T0 in pear (60AU / Conference, on the left) and apple (GalaRvi6 / Gala, on the right). The number of up- or down-regulated DEGs is expressed as a percentage of the total number of genes present in the pear V1 (49306 genes) and Ariane V2 (31311 genes) microarrays, respectively. DEGs are classified in functional categories according to MapMan 3.5.1R2 bins. Only bins with ≥ 6 DEGs are presented.

**Additional File 4: Table S5:** Metadata for the DEGs discussed in this work.

## Abbreviations

AUDPC: area under the disease progress curve
BR: brassinosteroid
CHS: chalcone synthase
CDPK: calcium dependent protein kinase
CRL: cullin ring ligase
DAMP: damage-associated molecular pattern
DEG: differentially expressed gene
DFR: dihydroflavonol 4-reductase
DGDG: digalactosyldiacylglycerol
ET: ethylene
ETI: effector triggered immunity
FA: fatty acid
hpi: hours post inoculation
FLS: flavonol synthase
HR: hypersensitive reaction
IAA: indole acetic acid
JA: jasmonic acid
KFB: Kelch motif containing F-box
LRR: leucine rich repeat
LTP: lipid transfer protein
MAMP: microbe-associated molecular pattern
MeJA: methyl jasmonate
MGDG: monogalactosyldiacylglycerol
nsLTP: non specific lipid transfer protein
NLR: nucleotide binding leucine-rich repeat
PAL: phenyl ammonia lyase
PCR: polymerase chain reaction
PME: pectin methyl esterase
PR: pathogenesis related
PRR: pattern recognition receptor
PTI: pattern triggered immunity
PUB: plant U-box type E3 ubiquitin
Q-PCR: quantitative polymerase chain reaction
QTL: quantitative trait locus
R: resistance
RLK: receptor like kinase
RLP: receptor like protein
RNA-seq: RNA sequencing
SA: salicylic acid
SAR: systemic acquired resistance
SCF: SKP1-CUL1-F box protein
SEM: scanning electronic microscopy
SKP: S phase kinase associated protein
TF: transcription factor
UGT: uridine 5’-diphospho-glucuronyl
VLCFA: very long chain fatty acid

## Declarations

## Acknowledgements

The authors gratefully acknowledge the IRHS-ImHorPhen team of INRA Angers for technical assistance in plant maintenance, B. Billy (SNES-GEVES) for technical assistance in flow cytometry and the technical platforms ANAN and IMAC.

## Authors contribution

EC, LP, EV and HS conceived the study. EC and EV supervised the study. ER, MB, PB and RC performed the biological experiments. SG and SP performed the database work and assisted with the bioinformatics analysis. LP wrote the original manuscript. EV, EC and HS edited the manuscript. All authors have read and agreed to the published version of the manuscript.

## Availability of data and materials

The datasets supporting the conclusion of this article are available in the Gene Expression Omnibus (GEO) repository [https://www.ncbi.nlm.nih.gov/geo/] with GSE159179 and GSE159180 accession numbers for apple and pear respectively.

## Ethics approval and consent to participate

This section is not applicable.

## Consent for publication

This section is not applicable.

