## Supplementary material for "Successful intergeneric transfer of a major apple scab resistance gene (*Rvi6*) from apple to pear and precise comparison of the downstream molecular mechanisms of this resistance in both species": Table S1 & Table S3

**Table S1**: Scab qualitative note of nine transgenic pear lines and non-transgenic Conference inoculated with three *V. pirina* strains. Percentage of plants in the different classes of symptoms, 42 days after inoculation.

| Class of symptoms | Conf. | 60C | 60S | 60W | 60AI | 60AK | 60AO | 60AS | 60AT | 60AU |
| --- | --- | --- | --- | --- | --- | --- | --- | --- | --- | --- |
| Strain VP102 | | | | | | | | | | |
| 0 | 0 | 0 | 0 | 0 | 0 | 0 | 0 | 0 | 0 | 0 |
| 1 | 0 | 0 | 0 | 0 | 0 | 0 | 0 | 0 | 0 | 0 |
| 2 | 0 | 40 | 41 | 12 | 20 | 56 | 56 | 88 | 24 | 23 |
| 3a | 0 | 60 | 59 | 50 | 60 | 44 | 40 | 12 | 60 | 77 |
| 3b | 0 | 0 | 0 | 0 | 20 | 0 | 0 | 0 | 4 | 0 |
| 4 | 100 | 0 | 0 | 38 | 0 | 0 | 0 | 0 | 12 | 0 |
| Strain VP132 | | | | | | | | | | |
| 0 | 0 | 0 | 0 | n.t. | 0 | n.t. | 0 | 0 | 0 | 0 |
| 1 | 0 | 0 | 0 | n.t. | 0 | n.t. | 0 | 0 | 0 | 0 |
| 2 | 0 | 75 | 58 | n.t. | 65 | n.t. | 67 | 53 | 36 | 37 |
| 3a | 0 | 25 | 42 | n.t. | 35 | n.t. | 33 | 47 | 56 | 63 |
| 3b | 0 | 0 | 0 | n.t. | 0 | n.t. | 0 | 0 | 8 | 0 |
| 4 | 100 | 0 | 0 | n.t. | 0 | n.t. | 0 | 0 | 0 | 0 |
| Strain VP98 | | | | | | | | | | |
| 0 | 0 | 0 | 0 | n.t. | 0 | n.t. | 0 | 0 | 0 | n.t. |
| 1 | 0 | 0 | 0 | n.t. | 0 | n.t. | 0 | 0 | 0 | n.t. |
| 2 | 0 | 31 | 10 | n.t. | 0 | n.t. | 9 | 7 | 5 | n.t. |
| 3a | 0 | 44 | 52 | n.t. | 33 | n.t. | 32 | 36 | 0 | n.t. |
| 3b | 0 | 13 | 33 | n.t. | 0 | n.t. | 27 | 14 | 10 | n.t. |
| 4 | 100 | 12 | 5 | n.t. | 67 | n.t. | 32 | 43 | 85 | n.t. |

Class 0: absence of symptoms

Class 1: hypersensitivity (pin points)

Class 2: resistance (chlorotic lesions, slight necrosis, crinkled aspect)

Class 3a: weak resistance (necrotic or chlorotic lesions with occasional very light sporulation)

Class 3b: weak susceptibility (clearly sporulating chlorotic or necrotic lesions

Class 4: susceptibility (sporulation only)

**Table S3**: Expression modulation of cell wall related DEGs detected at 8, 24 or 72 hpi during apple (Gala Rvi6 / Gala) and pear (60AU / Conference) responses to *V. pirina* and *V. inaequalis*, respectively. In red: up-regulated DEGs, in blue: down-regulated DEGs.

| Type of cell wall | Cell wall component | DEG name | Function | Expression modulation | |
| --- | --- | --- | --- | --- | --- |
|  |  |  |  | Apple | Pear |
| Primary | Cellulose | SHV3 | cellulose accumulation and pectin linking |  |  |
|  |  | SVL1 | cellulose accumulation and pectin linking |  |  |
|  |  | COB | glycosylphosphatidylinositol-anchored protein |  |  |
|  |  | CESA3 | biosynthesis |  |  |
|  |  | CESA6 | biosynthesis |  |  |
|  |  | CESA9 | biosynthesis |  |  |
| Primary | Hemicellulose: xyloglucans | MUR1 | xyloglucan galactosyltransferase |  |  |
|  |  | MUR2 | fucosyltransferase |  |  |
|  |  | MUR3 | GDP-L-fucose biosynthesis |  |  |
|  |  | XTH33 | xyloglucan/xyloglucosyl transferase |  |  |
|  |  | EXGT-A4 | xyloglucan/xyloglucosyl transferase |  |  |
|  |  | XTR2 | xyloglucan/xyloglucosyl transferase |  |  |
|  |  | XTR7 | loosening the network of cellulose and xyloglucan fibers |  |  |
|  |  | at5g20950 | beta-glucosidase involved in xyloglucan metabolism |  |  |
|  |  | at5g65730 | xyloglucan endotransglucosydase/hydrolase |  |  |
|  |  | at2g14620 | xyloglucan endotransglucosydase/hydrolase |  |  |
|  |  | at3g23730 | xyloglucan endotransglucosylase |  |  |
| Primary | Pectin | GAE1 | pectin biosynthesis |  |  |
|  |  | GAE3 | pectin biosynthesis |  |  |
|  |  | GAE6 | pectin biosynthesis |  |  |
|  |  | at3g16850 | pectinase |  |  |
|  |  | at3g42950 | pectin lyase-like |  |  |
|  |  | at4g23820 | pectin lyase-like |  |  |
|  |  | at3g61490 | pectin lyase-like |  |  |
|  |  | at5g20260 | pectin biosynthesis |  |  |
|  |  | GAUT1 | homogalacturonan biosynthesis |  |  |
|  |  | GAUT8 | homogalacturonan biosynthesis |  |  |
|  |  | at4g19420 | pectinacetylesterase |  |  |
|  |  | at5g26670 | pectinacetylesterase |  |  |
|  |  | at5g09760 | pectinesterase |  |  |
|  |  | at1g23200 | pectinesterase |  |  |
|  |  | ATPMEPCRF | pectin methylesterase PCR fragment F |  |  |
|  |  | ATPME3 | pectinesterase |  |  |
|  |  | PME31 | pectinesterase |  |  |
|  |  | RHM1 | rhamnose biosynthesis |  |  |
|  |  | NRS/ER | nucleotide-rhamnose synthase/epimerase-reductase |  |  |
|  |  | SHV3 | cellulose accumulation and pectin linking |  |  |
|  |  | SVL1 | cellulose accumulation and pectin linking |  |  |
|  |  | at1g70370 | degradation of homogalacturonan |  |  |
|  |  | at1g60590 | polygalacturonase |  |  |
|  |  | at2g43880 | polygalacturonase |  |  |
|  |  | at5g20260 | xylan and pectin biosyynthesis |  |  |
|  |  | AXS2 | rhamnogalacturonan II biosynthesis |  |  |
| Primary | Arabinogalactans | AGP1 | arabinogalactan protein |  |  |
|  |  | AGP18 | arabinogalactan protein |  |  |
|  |  | AGP20 | arabinogalactan protein |  |  |
|  |  | AGP26 | arabinogalactan protein |  |  |
|  |  | AGP17 | arabinogalactan protein |  |  |
|  |  | AGP14 | arabinogalactan protein |  |  |
|  |  | at3g22440 | hydroxyproline-rich glycoprotein family protein |  |  |
|  |  | at3g19020 | leucine-rich repeat family protein (related to HRPG) |  |  |
| Secondary | Cellulose | ATCSLC12 | biosynthesis |  |  |
| Secondary | Hemicellulose: xylans | DET3 | modification of level of wall-bound xylose |  |  |
|  |  | IRX6/COBL4 | modification of level of wall-bound xylose |  |  |
|  |  | IRX9 | family 43 glycosyl transferase |  |  |
|  |  | PGSIP3 | glucuronyltransferase |  |  |
|  |  | KOR1/IRX2 | biosynthesis |  |  |
|  |  | at5g20260 | xylan and pectin biosynthesis |  |  |
| Secondary | Lignin | C4H | biosynthesis |  |  |
|  |  | at5g14700 | biosynthesis |  |  |
|  |  | HCT | cinnamoyl-CoA reductase-related |  |  |
|  |  | OMT1 |  |  |  |
|  |  | 4CL3 |  |  |  |
|  |  | at2g23910 |  |  |  |
|  |  | C4H | cinnamoyl-CoA reductase-related |  |  |
|  |  | PAL1 |  |  |  |
|  |  | PAL2 |  |  |  |
|  |  | UGT72E1 |  |  |  |
|  |  | LAC6 |  |  |  |
