## Supplementary figures and images for "Successful intergeneric transfer of a major apple scab resistance gene (*Rvi6*) from apple to pear and precise comparison of the downstream molecular mechanisms of this resistance in both species"

### Fig. S1

#### Slide 1
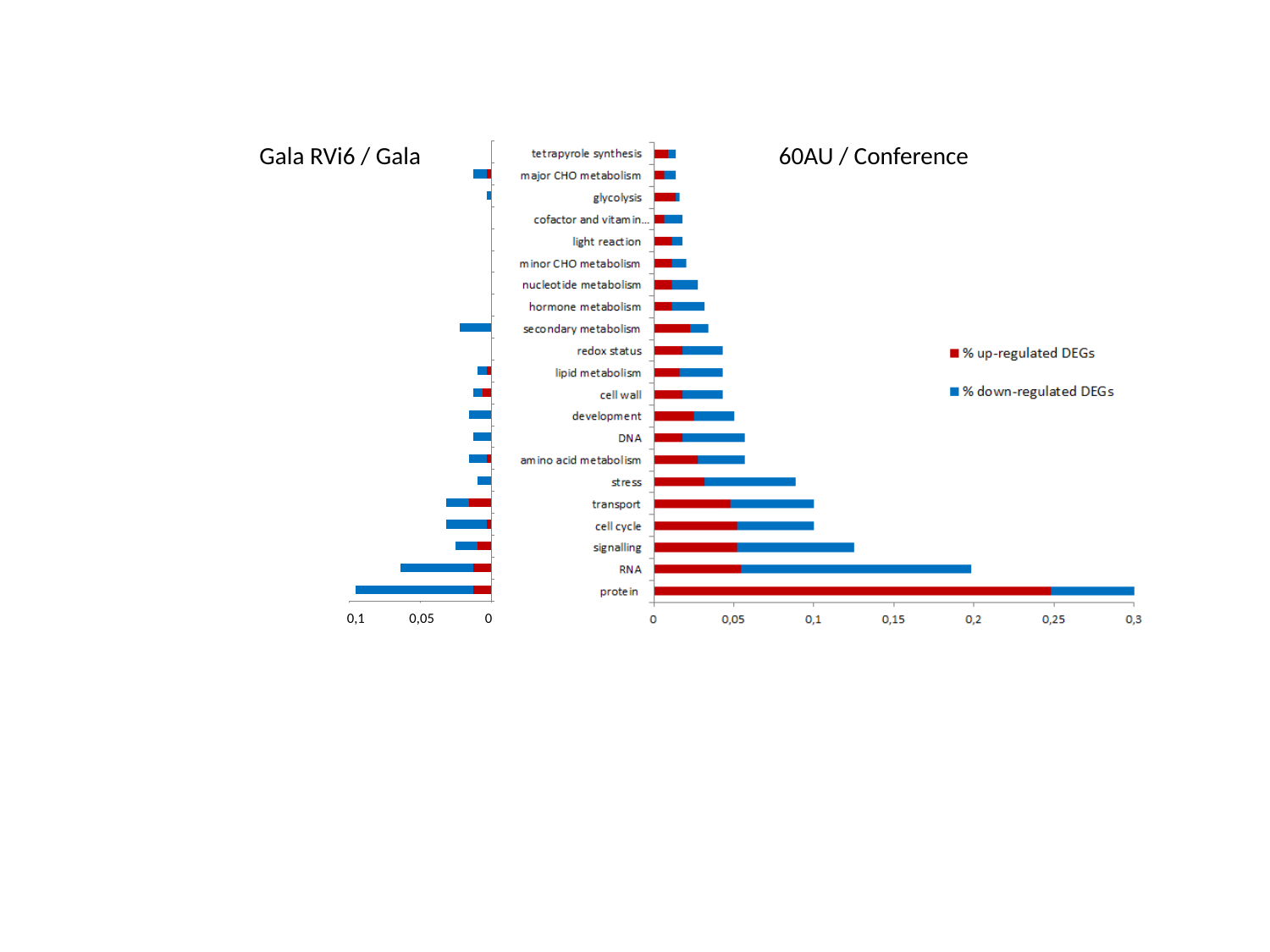

Gala RVi6 / Gala
60AU / Conference
 0,1 0,05 0
